# A Dysregulated Peripheral Immune Landscape Favors the Expansion of Epstein-Barr Virus-positive Age-Associated B Cells in Multiple Sclerosis

**DOI:** 10.64898/2026.11.16.767068

**Authors:** Leena Yoon, Sun Sook Chung, Simon Thebault, Gautier Breville, Leonardo Josué Castro Muñoz, Lauren N. MacMullen, Alina Gu, Tyler H. N. Yang, Songtao He, Daniel Claiborne, Avi Srivastava, Rena R. Xian, Richard F. Ambinder, Amit Bar-Or, Samantha S. Soldan, Paul M. Lieberman

## Abstract

Epstein-Barr virus (EBV) is a major risk factor for multiple sclerosis (MS), but the underlying mechanisms remain poorly understood. Here, we use single-cell RNA sequencing coupled with targeted EBV transcript enrichment to characterize host immune and viral transcriptional dynamics across multiple time points following EBV infection of PBMCs from patients with active MS compared to healthy controls (HC). Prior to *ex vivo* infection, PBMCs from MS patients exhibited distinct immune cell composition and transcriptional profiles, including increased detection of EBV transcripts in monocytes. Early responses to EBV were dominated by changes in monocyte and NK cells populations and transcript states. At later time points, MS-associated B- and T-cell responses dominated, characterized by the expansion of EBV-infected age-associated B-cells (ABCs) and increased expression of T-cell exhaustion-associated markers. MS-associated EBV-infected ABCs were deregulated for EBV lytic genes and cellular pathways associated with interferon-alpha responses, antigen processing and presentation, and Toll-like receptor signaling. Together, these findings define an MS-associated inflammatory immune state and dysfunctional response that favors the formation of pathogenic EBV-infected ABCs.

## Introduction

Multiple Sclerosis (MS) is a chronic autoimmune disease of the central nervous system (CNS) affecting over 3.1 million individuals worldwide^1-3^. MS is a heterogeneous disease characterized by complex interactions between genetic susceptibility and environmental risk factors^2,3^. Numerous studies have identified Epstein-Barr virus (EBV) as the strongest established environmental risk factor with compelling epidemiological and mechanistic evidence supporting a causal role in MS pathogenesis^4-6^.

EBV is a ubiquitous human gamma herpesvirus that infects more than 95% of the adult population and establishes lifelong persistent infection^7-10^. Primary EBV infection is the major cause of infectious mononucleosis (IM), and although most infections are asymptomatic or resolve without long-term complications, EBV is associated with diverse malignancies, including B-cell lymphomas, T/NK cell lymphomas, nasopharyngeal and gastric carcinomas, and rare leiomyosarcomas^11,12^. Recent studies have provided strong evidence for a role of EBV in various autoimmune disorders, including rheumatoid arthritis, systemic lupus erythematosus (SLE) and MS^12,13^. In healthy individuals, EBV persists predominantly as a latent infection in long-lived resting memory B cells, but sporadic lytic cycle reactivation occurs in the oropharynx through poorly understood mechanisms^8,14,15^. The balance between viral latency and lytic reactivation is controlled largely through EBV-specific cellular and humoral immunity^16,17^. Most EBV-associated disease is linked to imbalances in this EBV-specific immunity, suggesting that altered virus-host dynamics are involved in MS pathogenesis.

Several non-mutually exclusive mechanisms have been proposed to explain how EBV may contribute to MS, including impaired immune control of EBV-infected cells that can lead to aberrant B- and T-cell activation, autoantigen presentation and CNS trafficking^5,6,18^. EBV infection has been shown to induce T-bet^+^CXCR3^+^ B cells with enhanced CNS-homing properties^19^, while related TBX21^+^CD21^lo^CXCR3^+^ age-associated B-cell (ABC) populations are expanded in early MS and associated with disease activity^20,21^. EBV infection can also alter B-cell antigen processing and presentation, resulting in HLA-DR15 presentation of myelin basic protein (MBP) peptides that were also identified in MS brain tissue and recognized by peripheral and CSF-derived CD4^+^ T cells from patients with MS^22^. More recently, longitudinal analysis of patients with relapsing-remitting MS identified EBV-associated transcriptional changes in B cells and monocytes before clinical relapse, accompanied by expansion of EBV-infected ABC-like B-cell populations^23^. Moreover, EBV transcription factors, especially EBNA2, preferentially interact with host regulatory regions containing genetic variants associated with B-cell activation in MS and other autoimmune diseases. These observations suggest a potential disease-driving mechanism where viral infection may interact with host genetic risk^24^. Studies from our lab found that spontaneously growing EBV positive B cell lines, known as spontaneous lymphoblastoid lines (SLCLs), were generated more frequently from patients with MS and exhibited increased expression of lytic EBV genes together with altered interferon and inflammatory host responses and EBV-host interactions in MS^25^. Collectively, these studies suggest that differences in genetic susceptibility, immune regulation, and B-cell control of EBV latency may contribute to MS pathogenesis.

The cellular mechanisms underlying altered immune responses to EBV infection in MS remain incompletely understood. Previous studies examining EBV in MS have analyzed patient-derived immune cells, isolated specific immune cell populations, or established EBV-positive cell lines. Although these approaches have provided important insights into EBV-associated immune responses in MS, they do not consider how innate and adaptive immune cells influence EBV infection outcomes and how this may be altered in MS. An *ex vivo* peripheral blood mononuclear cell (PBMC) infection model provides an opportunity to address these questions while maintaining interactions among the major immune populations present in peripheral blood. We previously established an *ex vivo* PBMC model to characterize the cellular and transcriptional responses to EBV infection in healthy adults^26^. Longitudinal immune profiling revealed that the earliest response to EBV was dominated by monocytes, followed by T-cell and NK-cell responses associated with subsequent B-cell outgrowth. Targeted enrichment of EBV transcripts further enabled characterization of viral gene expression to be followed during the transition from early infection to B-cell transformation and latency. These findings demonstrated that EBV infection outcomes are influenced by immune populations outside the B-cell compartment, which vary significantly among individuals. As most adults have been previously exposed to EBV and harbor latent infection, this model allows us to compare immune responses to a standardized EBV exposure across individuals with different preexisting immune states. By using the same viral inoculum, we can examine differences in host immune responses and EBV infection outcomes while maintaining interactions among peripheral immune populations.

Here, we used this *ex vivo* PBMC infection model to investigate how preexisting peripheral immune states and subsequent responses to EBV differ between patients with MS and healthy controls. We combined longitudinal single-cell RNA sequencing based on split-pool combinatorial barcoding and targeted EBV transcript enrichment with flow cytometry and bulk RNA sequencing to compare the response to *ex vivo* EBV infection in PBMCs from patients with MS and healthy controls. PBMCs were exposed to the same EBV inoculum and analyzed at Days 1, 7, and 15, allowing host transcriptional responses and EBV gene expression to be followed across myeloid, NK-, T-, and B-cell populations. We identified MS-specific differences in cell composition and transcriptomes for monocytes, NK, T and B-cell states, some of which were present before EBV exposure. We identified EBV transcripts predominantly in newly infected B-cells, but also in monocytes at early time points and prior to ex vivo infection. T-cell exhaustion-related markers and expansion of EBV-infected atypical/age-associated B-cell (ABC) populations were more prevalent in active MS than in healthy controls. Viral and host transcriptional signatures could be defined for EBV-infected ABCs which are implicated in MS pathogenesis. Together, these findings identify MS-specific differences in peripheral immune states and a distinct temporal progression of innate and cellular immune responses to EBV consistent with known immunopathogenic features in MS.

## RESULTS

### MS patient PBMCs are more permissive to EBV infection and B-cell immortalization

PBMCs from individuals with MS (n = 19), non-MS neurological controls (n = 5), and healthy controls (HC; n = 15) were infected with EBV strain Mutu I at a multiplicity of infection (MOI) of 1 or maintained under matched mock conditions (Figure 1A) in the absence of cyclosporin A, which is routinely used to eliminate T cells from PBMC cultures to enable the emergence of EBV-transformed lymphoblastoid lines (LCLs). Cellular and transcriptional responses were characterized by flow cytometry, bulk RNA-seq, and single-cell RNA sequencing longitudinally across timepoints. For bulk RNA-seq, PBMCs from 12 MS donors, five non-MS neurological controls, and five HC donors were analyzed at Days 1 and 7 following *ex vivo* EBV infection (Supplementary Figure S1A). For single-cell analysis, PBMCs from eight MS donors, including five patients with MRI-defined active MS and three patients with stable MS, together with four HC donors, were analyzed at Days 1, 7, and 15 following *ex* vivo EBV infection (Table S1). The bulk and single-cell RNA-seq cohorts included one overlapping donor. All donors tested were anti-EBNA1 IgG seropositive, suggesting that these assays measured response to re-exposure of EBV.

**Figure 1.**
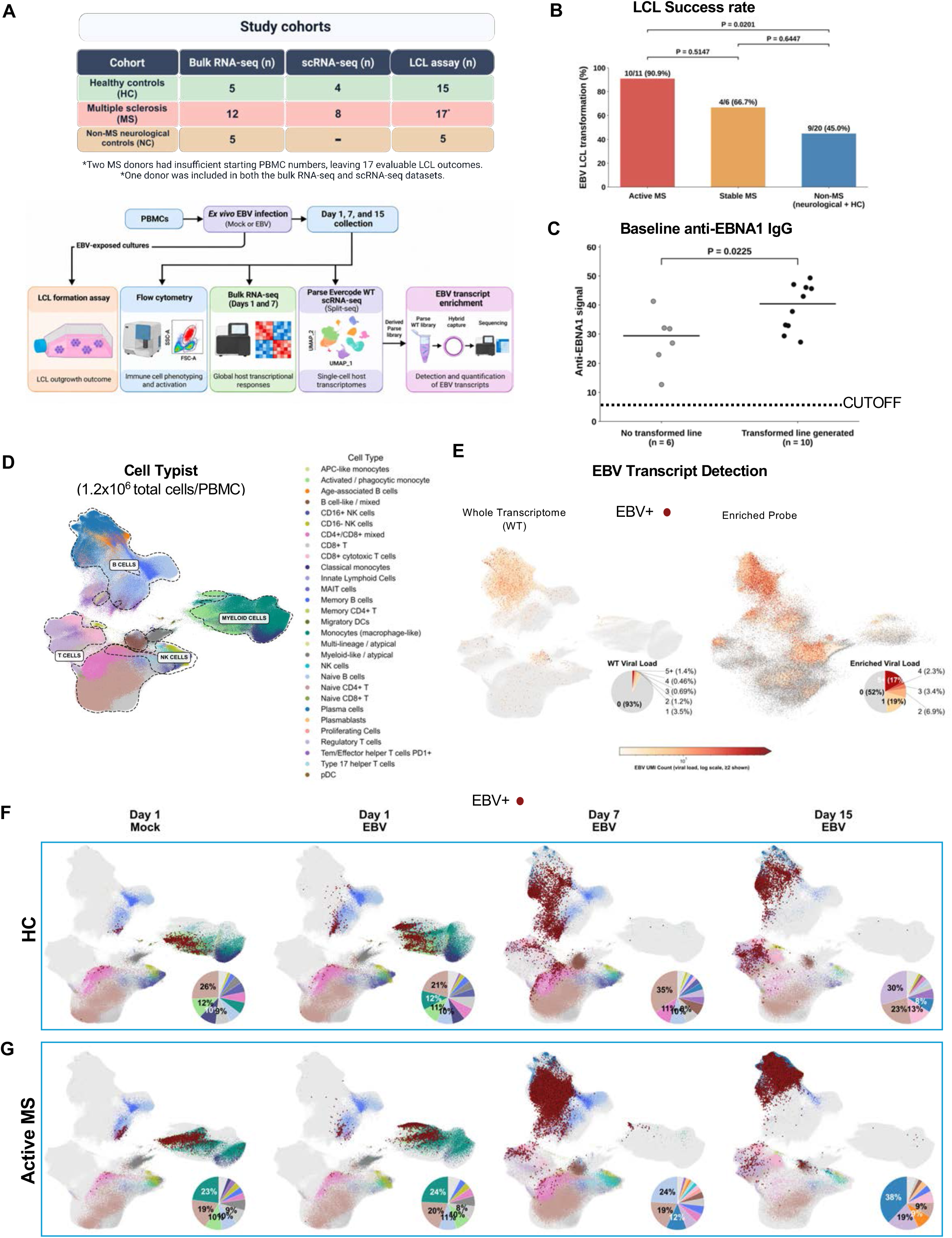
Bulk and Single-cell profiling of immune-cell responses and EBV transcript distribution following *ex vivo* infection of PBMCs. (A) Study design and experimental workflow. PBMCs from healthy controls (HC), individuals with multiple sclerosis (MS), and non-MS neurological controls were cultured under mock or EBV-exposed conditions and collected at the indicated time points for flow cytometry, bulk RNA sequencing, and single-cell whole-transcriptome profiling with targeted EBV transcript enrichment. Cohort sizes for each experimental analysis are indicated. (B) Frequency of lymphoblastoid cell line (LCL) formation following ex vivo EBV infection in Active MS (10/11, 90.9%), Stable MS (4/6, 66.7%), and non-MS controls (9/20, 45.0%; neurological controls and HC). Two MS donors were excluded because of insufficient starting PBMC numbers. Statistical significance was assessed using two-sided Fisher’s exact tests. (C) Baseline anti-EBNA antibody signals in donors with (n = 10) or without (n = 6) successful EBV-transformed cell-line generation. Only donors with available anti-EBNA measurements and evaluable transformation outcomes were included. Horizontal bars indicate group means. Statistical significance was assessed using a two-sided Mann–Whitney U test. (D) UMAP embedding of approximately 1.29 million PBMCs, colored by cell type. CellTypist annotations were refined using canonical marker gene expression. Dashed contours indicate major immune-cell lineages. (E) Comparison of EBV transcript detection using conventional whole-transcriptome (WT) single-cell RNA sequencing and targeted EBV transcript enrichment. Cells are colored by EBV-derived UMI count, and inset pie charts show the distribution of viral UMI counts per cell. (F, G) UMAP projections showing immune-cell populations and EBV-positive cells in HC (F) and Active MS (G) under mock conditions and following EBV exposure on days 1, 7, and 15. EBV-positive cells are overlaid in dark red. Inset pie charts indicate the cell-type composition of EBV-positive cells. EBV-positive cells were defined as cells with ≥2 EBV-derived UMIs spanning at least two distinct EBV genes.

Differences between groups were first identified in Lymphoblastoid cell lines (LCLs) formation following EBV infection. LCLs were generated more frequently from active MS PBMC cultures (90.9%, 10/11) than from stable MS (66.7%, 4/6) or non-MS cultures (45.0%, 9/20) (Figure 1B). The difference between active MS and non-MS was nominally significant (two-sided Fisher’s exact test, P = 0.0201). Two MS donors were excluded because of insufficient starting PBMC numbers. Clinical data were available for most of the patient samples tested for LCL formation (Supplemental Table S1). Among the clinical characteristics measured, only EBNA1 antibody levels were significantly associated with LCL outcomes (Figure. 1B). Baseline anti-EBNA signals were higher in those whose cultures generated EBV-transformed cell lines (n = 10) compared with those without successful transformation (n = 6; P = 0.0225, two-sided Mann–Whitney U test; Figure 1C). No significant associations were identified for other measured clinical variables, including EBV DNA levels assessed by ddPCR, neurofilament light chain (NfL) levels, or CSF IgG index. These findings suggest that preexisting immune responses to EBV may influence subsequent EBV-driven B-cell outgrowth.

To characterize immune responses to *ex vivo* EBV infection, we first used bulk RNA-seq on PBMCs from 22 donors to examine differences in the transcriptional response to EBV between MS and non-MS PBMCs (Supplementary Figure S1B–D). Principal component analysis showed separation according to time following EBV exposure, with differences between MS and non-MS samples also observed at each time point (Supplementary Figure S1B). Differential gene expression and pathway analysis further identified distinct transcriptional programs between MS and non-MS PBMCs following EBV *ex vivo* infection (Supplementary Figure S1C). To examine whether peripheral transcriptional responses to EBV were associated with clinical characteristics of MS, we integrated donor-matched bulk RNA-seq data with clinical measurements (Supplementary Figure S1E–F). Day 1 EBV-induced B-cell differentiation-associated transcriptional responses were inversely correlated with CSF IgG index (Spearman ρ = −0.76, P = 0.004, n = 12). These exploratory findings suggested associations between peripheral EBV-induced transcriptional responses and measures of intrathecal humoral immune activity, although neither association remained significant after correction for multiple comparisons.

To better understand immune cell population dynamics in response to *ex vivo* EBV infection, we performed single-cell RNA-seq using the high-throughput Parse Biosciences split-pool combinatorial barcoding platform. Following quality control, the combined dataset contained approximately 1.29 million cells comprising 29 annotated immune cell populations across the major B-cell, T-cell, NK-cell, and myeloid lineages (Figure 1D). Initial cell-type annotations were assigned using CellTypist and subsequently validated using canonical lineage and subtype marker expression (Supplementary Figure S2). To overcome the limited sensitivity of conventional single-cell RNA sequencing for EBV transcript detection, we used an EBV genome-wide probe set for targeted hybridization capture (Fig. 1D). This approach substantially improved viral RNA detection compared with conventional single-cell RNA sequencing without viral enrichment. The proportion of cells with at least two detectable EBV-derived UMIs increased from 3.8% in WT libraries to 29.4% in EBV-enriched libraries, accompanied by an increase in the number of viral transcripts detected per cell (Figure 1E). Mapping EBV-positive cells onto the global WT landscape showed a marked change in their cellular distribution over time (Figure 1F–G). At Day 1, EBV-positive cells were detected across multiple immune populations, with a greater contribution from myeloid populations in MS compared with HC. At later time points, EBV-positive cells shifted toward lymphoid populations and became increasingly concentrated within the B-cell compartment by Day 15. This transition was more pronounced in MS, with greater accumulation of EBV-positive B cells, consistent with increased frequency of LCL formation.

### MS-specific immune cell population differences at baseline and in response to EBV infection

We next assessed the differences in immune cell populations between MS and HC in response to *ex vivo* EBV infection. EBV-positive cells were quantified across immune populations and time points (Figure 2A–B). At Day 1, differences were most apparent within the myeloid compartment, including baseline mock-infected samples. Activated/phagocytic monocytes showed higher EBV positivity in active MS than HC (13.3% versus 5.7%), and this difference increased by Day 7 (26.7% versus 12.3%). Differences were also observed in macrophage-like monocytes at Day 1. In contrast, EBV positivity remained low in classical monocytes, indicating that the early myeloid-associated viral signal was concentrated within specific activated monocyte states rather than uniformly distributed across the monocyte compartment. At later time points, the largest differences were observed in differentiated B-cell populations. By Day 15, EBV positivity reached 63.2% in plasmablasts and 54.7% in Age-Associated B-cells (ABCs) from active MS donors, compared with 33.7% and 26.3% in HC, respectively. Plasma cells also showed higher EBV positivity in active MS (37.6% versus 18.1%), whereas memory B cells showed a moderate difference (23.2% versus 18.8%) (Figure 2A).

**Figure 2.**
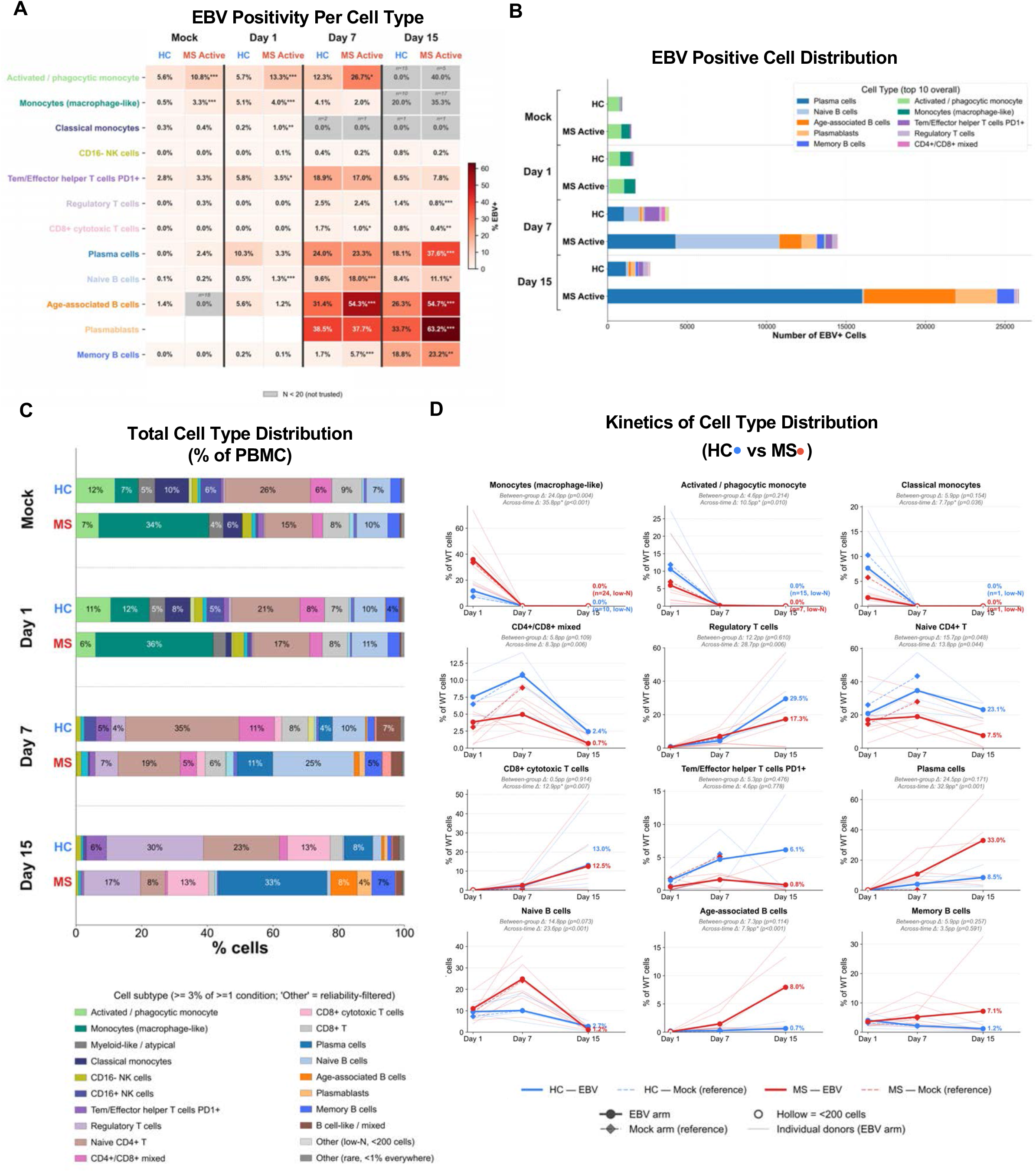
Temporal changes in EBV positivity and immune-cell composition following EBV infection. (A) Heatmap showing the percentage of EBV-positive cells within each annotated cell type in healthy controls (HC) and Active MS samples under mock conditions on day 1 and following EBV exposure on days 1, 7, and 15. EBV-positive cells were defined as cells with ≥2 EBV-derived UMIs spanning at least two distinct EBV genes. Gray tiles indicate populations containing fewer than 20 cells. Asterisks denote Fisher’s exact test comparisons between active MS and HC within the same timepoint (* p<0.05, ** p<0.01, *** p<0.001). (B) Absolute numbers of EBV-positive cells by cell type, displayed using the same Disease Group (HC, stable MS, active MS) × Timepoint (Mock, Day 1, Day 7, Day 15) layout as in (C). Bars are stacked and colored by cell type. Legend includes the 10 cell types contributing the largest number of EBV-positive cells across the full-time course. (C) Relative proportions of annotated cell types among total PBMCs across groups and conditions. Cell types representing ≥3% of cells in at least one condition are displayed individually; remaining populations are grouped as indicated. (D) Donor-level kinetics of selected cell populations following EBV exposure. Blue and red indicate HC and MS, respectively. Thin lines represent individual donors, and bold solid lines show group trajectories. Dashed lines indicate mock reference trajectories. Open symbols indicate observations containing fewer than 200 cells. Panel subtitles summarize the maximum between-group difference (HC versus MS at the time point showing the largest percentage-point separation; two-sided exact Mann-Whitney U test) and the maximum within-group temporal change (largest Day 1–Day 15 trajectory shift; donor-blocked permutation test, 2,000 permutations).

The absolute number of EBV-positive cells showed a similar increase over time (Figure 2B). Relatively few EBV-positive cells were detected at Day 1, followed by a marked increase at Days 7 and 15. This increase was substantially greater in active MS, particularly at Day 15, and was dominated by B-cell populations. In contrast, PBMCs from stable MS patients showed a smaller increase in EBV-positive cells and did not reach the magnitude observed in active MS (Supplementary Figures S3). Donor-level analysis of the single-cell data, normalized to equal cell numbers, similarly showed higher EBV transcript abundance in active MS than HC at Days 7 and 15 (Supplementary Figure S4A), indicating that the increased EBV signal was observed across donors and was not solely driven by differences in the number of cells analyzed.

The overall immune cell composition also changed substantially over the infection time course (Figure 2C–D). Myeloid populations decreased after the early time points, while NK-, T-, and B-cell populations showed different patterns at later time points. Cytotoxic NK cells (CD16^+^ NK cells) were lower in MS at baseline, while several T-cell populations including regulatory T cells, PD1^+^ Tem/effector helper T cells, and cytotoxic CD8^+^ T cells, also showed different trajectories between MS and HC. Within the B-cell compartment, naive B cells initially increased and subsequently declined, whereas ABCs and plasma cells increased at later time points, with greater changes observed in MS (Figure 2D). Together, these findings showed that the differences between MS and HC existed prior to infection, and further diverged over the course of EBV infection, beginning with differences in myeloid and NK-cell populations and followed by changes in T-cell and differentiated B-cell populations at later time points.

### MS monocytes show pre-existing transcriptional differences and enhanced antiviral responses to EBV

The early divergence in the myeloid compartment prompted a more detailed analysis of monocyte states. Transcriptional annotation resolved classical, activated/phagocytic, macrophage-like, APC-like, atypical, and additional myeloid populations (Figure 3A). The distribution of these populations differed between MS and HC, with macrophage-like monocytes representing a larger proportion of the monocyte compartment in MS (Figure 3A, Supplementary Figure S5A). Differences were also present before *ex vivo* infection. At baseline (Mock Day 1), monocytes from active MS donors showed increased expression of genes associated with immune and inflammatory responses, lipid metabolism, cell adhesion and migration, and cellular differentiation compared with HC (Figure 3B). Lipid metabolism genes, including PLPP3, FABP4, PLIN2, APOC1, APOE, and FABP5, were consistently higher in active MS, together with inflammatory genes including LEP, CCL18, GSDME, HAMP, and TFRC.

**Figure 3.**
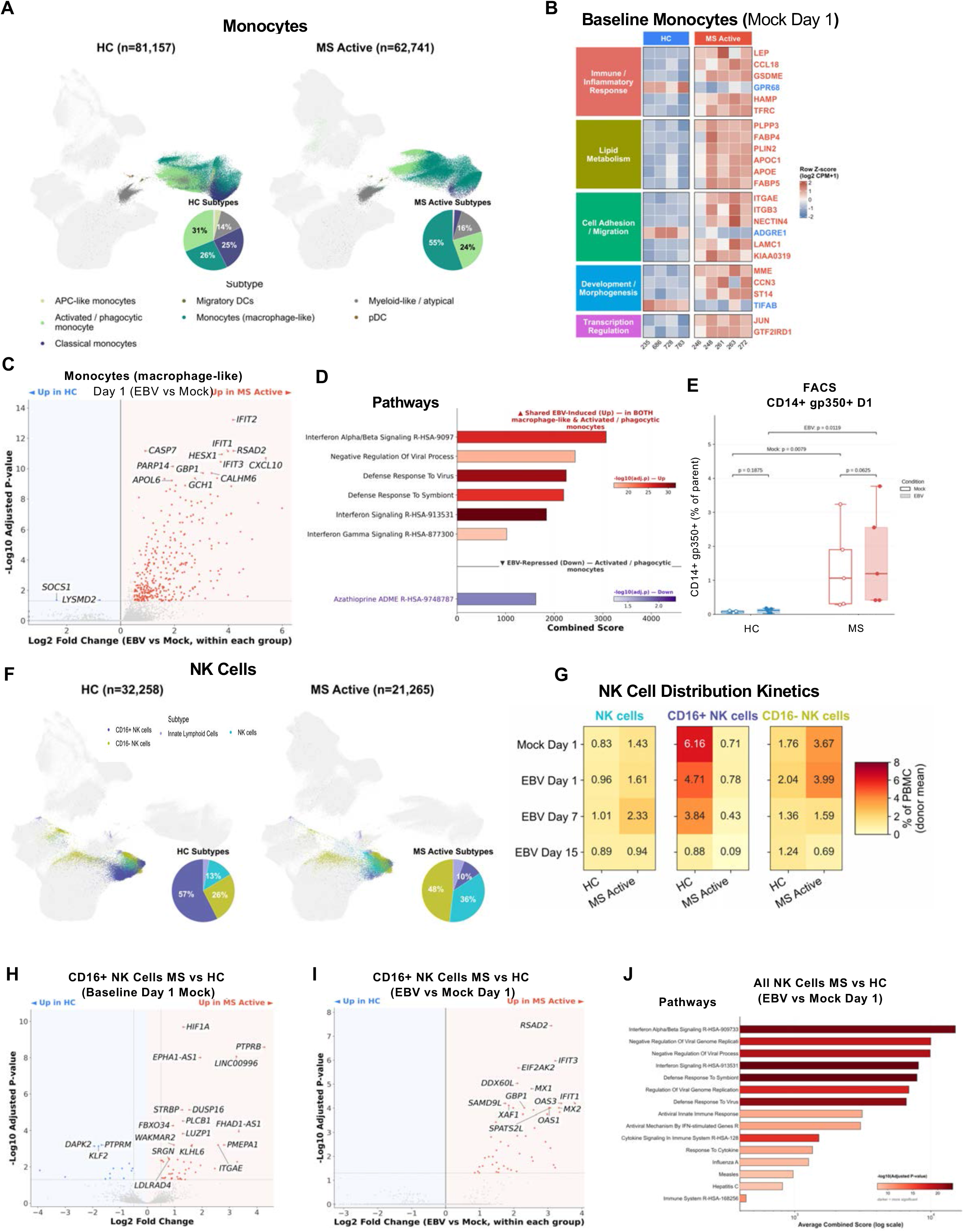
Monocytes and NK cells from active MS donors showed baseline transcriptional differences and amplified interferon responses following EBV infection. (A) UMAP plots showing monocyte and related myeloid populations in healthy controls (HC) and active MS donors, colored by annotated subtype. Inset pie charts indicate subtype proportions. (B) Row Z-scored pseudobulk expression heatmap of genes differentially expressed between active MS and HC monocytes under baseline mock conditions on day 1. Combined monocytes comprise classical, activated/phagocytic, and macrophage-like monocytes. Columns represent donors, and genes are grouped by functional category. (C) Chimeric volcano plot showing paired day 1 EBV-versus-mock transcriptional responses in macrophage-like monocytes, analyzed separately within HC (left) and active MS (right). (D) Pathways enriched among EBV-induced genes shared by macrophage-like and activated/phagocytic monocytes from active MS donors. Bar length indicates the mean combined enrichment score across both subtypes, and color indicates −log₁₀(adjusted P value). The lower bar shows a pathway enriched among EBV-repressed genes in activated/phagocytic monocytes. (E) Flow cytometric quantification of CD14⁺gp350⁺ cells in mock- and EBV-exposed samples from HC and MS donors on day 1 (n = 5 donors per group). Points represent individual donors. Statistical significance was assessed using two-sided Wilcoxon matched-pairs signed-rank tests for mock-versus-EBV comparisons within groups and two-sided Mann–Whitney U tests for comparisons between HC and active MS donors. (F) UMAP plots showing NK-cell and related populations in HC and active MS donors, colored by annotated subtype. Inset pie charts indicate subtype proportions. (G) Heatmaps showing donor-mean frequencies of total NK cells, CD16⁺ NK cells, and CD16⁻ NK cells as percentages of PBMCs across the indicated conditions and time points. (H) Baseline differential gene expression between active MS and HC CD16⁺ NK cells under mock conditions on day 1. (I) Chimeric volcano plot showing day 1 EBV-versus-mock transcriptional responses in CD16⁺ NK cells, analyzed separately within HC (left) and active MS (right). (J) Pathways significantly enriched among EBV-induced genes in all three NK-cell populations from active MS donors on day 1, ranked by average combined enrichment score. Color indicates −log₁₀(adjusted P value).

Following EBV infection, macrophage-like monocytes from active MS donors showed a broader transcriptional response at Day 1 compared with HC (Figure 3C). A similar pattern was observed in activated/phagocytic monocytes (Supplementary Figure S5B). Despite differences in the number and magnitude of differentially expressed genes between the two monocyte populations, pathway analysis identified a common response in active MS, including interferon alpha/beta and interferon gamma signaling, defense response to virus, and regulation of viral processes (Figure 3D). Consistent with transcriptional differences in inflammatory and cell-migration programs, flow cytometry revealed that CCR1- and CCR2-expressing monocyte populations also responded differently to EBV. CCR1^+^ CCR2^+^ and CCR2^+^ monocytes decreased following EBV exposure in HC but were maintained in MS (Supplementary Figure S6C–E).

We also identified EBV-positive cells within the myeloid compartment at early time points in both mock and EBV infected PBMC (Figures 1F–G and 2A). EBV positivity was particularly enriched within activated/phagocytic and macrophage-like monocytes, while remaining relatively low in classical monocytes. Viral gene expression profiles also differed between active MS and HC, with differences in the frequency of cells expressing individual EBV genes and their expression levels across myeloid populations (Supplementary Figure S5C). To determine whether the early myeloid-associated EBV signal could also be detected at the protein level, we examined surface gp350 expression on CD14^+^ monocytes by flow cytometry. Notably, surface gp350 may reflect bound virions or phagocytosed material rather than productive monocyte infection. CD14⁺gp350⁺ cells were more frequent in MS than HC following *ex vivo* EBV infection at Day 1 (P = 0.0119; Figure 3E). Representative flow cytometry and the monocyte gating strategy are shown in Supplementary Figure S6A–B.

Together, these results showed that monocytes from MS donors were transcriptionally different from HC at baseline and developed a stronger antiviral response following infection. This response was concentrated within activated/phagocytic and macrophage-like monocyte states and was accompanied by increased detection of EBV RNA and gp350 at the early time point.

### NK cells in active MS show reduced CD16^+^ populations but enhanced early antiviral responses

Differences in the early response extended beyond myeloid cells. NK cells were separated into CD16^+^, CD16^−^, and additional NK-cell populations (Figure 3F). The frequency of the CD16^+^ NK cell population was markedly lower in active MS than HC at baseline (Mock Day 1) (Figure 3G). Following EBV infection, CD16^+^ NK cells progressively decreased over time in both groups, with a more pronounced decline in HC, which had a substantially higher baseline frequency. In active MS, CD16^+^ NK cells were already infrequent at baseline and declined further at later time points. PBMCs from donors with stable MS showed an intermediate NK-cell distribution, with low representation of CD16^+^ NK cells like active MS (Supplementary Figure S7A–B).

Transcriptional responses of NK cells from active MS donors were significantly different from those of HCs. At baseline (Mock Day1), active MS and HC differed in genes associated with signaling, cell adhesion, and cellular metabolism (Figure 3H). Pathway analysis identified differences in MAPK-associated signaling, phosphatase activity, cell-matrix adhesion, and related regulatory pathways (Supplementary Figure S7C). Following EBV infection, CD16^+^ NK cells showed increased induction of interferon-responsive and antiviral genes, including RSAD2, IFIT3, EIF2AK2, DDX60L, MX1, XAF1, OAS1, and OAS3 (Figure 3I), particularly in active MS. A similar response was observed in CD16^−^ NK cells at Day 1 (Supplementary Figure S7D). Pathway analysis further identified shared EBV-induced responses across NK-cell populations in active MS, including interferon signaling, defense responses to virus, and regulation of viral replication (Figure 3J; Supplementary Figure S7E). By Day 7, antiviral gene induction was observed in both active MS and HC across CD16+ and CD16^−^ NK-cell populations (Supplementary Figure S7F–G). Pathway analysis identified substantial overlap in interferon signaling, defense responses to virus, and regulation of viral replication across both groups (Supplementary Figure S7I–J). These results showed that NK cells from active MS had a more prominent antiviral transcriptional response at Day 1, whereas HC showed substantial induction of overlapping antiviral genes and pathways by Day 7.

Targeted viral enrichment also detected EBV transcripts in CD16^−^ NK cells (Supplementary Figure S7H). In contrast, no EBV transcripts were detected in CD16^+^ NK cells, despite the strong antiviral transcriptional response in this population. These findings distinguished the antiviral response of CD16^+^ NK cells from direct detection of viral transcripts within the same population.

### EBV induces distinct cytotoxic and activation programs across T-cell subsets in active MS

By Day 7, differences extended into the adaptive immune compartment. Transcriptional annotation identified multiple CD4^+^ and CD8^+^ T-cell populations, including naïve and memory CD4^+^ cells, regulatory T cells, PD1^+^ effector/memory helper T cells, CD8^+^ populations, cytotoxic CD8^+^ T cells, Th17 cells and MAIT cells (Figure 4A). PBMCs from donors with stable MS showed a similar overall distribution of T-cell populations by Day 7 (Supplementary Figure S8A).

**Figure 4.**
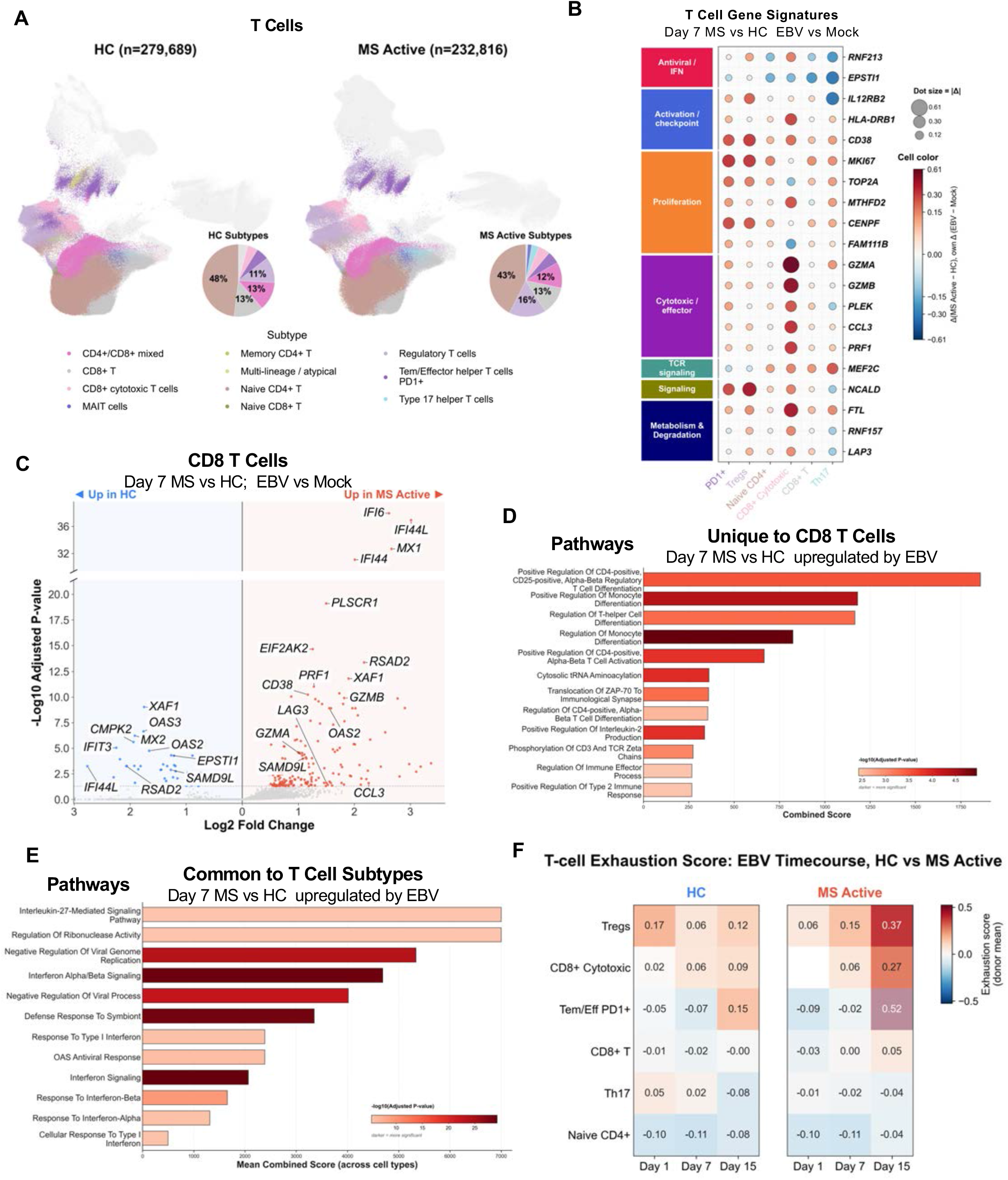
T-cell subsets show distinct antiviral, cytotoxic, and exhaustion-associated responses to EBV challenge in active MS donors. (A) UMAP embeddings showing T-cell populations in healthy controls (HC) and active MS donors, colored by annotated subtype. Inset pie charts indicate subtype proportions. (B) Subtype-specific differences in transcriptional responses to EBV infection on day 7. Values represent the difference in log₂ fold change for EBV versus mock treatment between Active MS and HC donors. Genes were selected based on the largest cross-subtype range of this difference across the six T-cell populations shown. Red indicates a greater response in active MS, and blue indicates a greater response in HC. Dot size represents the absolute difference, and color intensity reflects the magnitude and direction of the difference. Selected genes are grouped by functional category. (C) Chimeric volcano plot showing day 7 EBV-versus-mock transcriptional responses in CD8⁺ cytotoxic T cells, analyzed separately within HC (left) and active MS (right). The y-axis indicates −log₁₀(adjusted P value), with an axis break as shown. (D) Pathways uniquely enriched among EBV-upregulated genes in CD8⁺ cytotoxic T cells from active MS donors, relative to PD1⁺ Tem/effector helper T cells and regulatory T cells. Bar length indicates the combined enrichment score, and color indicates −log₁₀(adjusted P value). (E) Pathways enriched among EBV-upregulated genes in all three T-cell populations from active MS donors. Bar length indicates the mean combined enrichment score across the three populations, and color indicates −log₁₀(adjusted P value). (F) Heatmaps showing donor-mean T-cell exhaustion signature scores across the indicated T-cell subsets on days 1, 7, and 15 following EBV exposure in HC and active MS donors.

To identify T-cell responses that differed between active MS and HC, we compared the Day 7 EBV-induced responses (relative to Mock) across T-cell populations (Figure 4B). Differences in EBV-induced log₂ fold changes were observed between active MS and HC across antiviral/interferon, activation, proliferation, and cytotoxic/effector gene programs. PD1^+^ Tem/effector helper T cells and Tregs showed greater induction of activation and proliferation-associated genes, including CD38, MKI67, and CENPF, in active MS. The strongest differences were observed in CD8^+^ cytotoxic T cells, with greater induction of cytotoxic genes (GZMA, GZMB, and PRF1) and the inflammatory chemokine CCL3 in active MS compared with HC. In contrast, several antiviral genes, including RNF213 and EPSTI1, showed greater EBV-induced responses in HC across multiple T-cell populations.

In CD8^+^ cytotoxic T cells, active MS showed greater induction of cytotoxic and effector-associated genes, including GZMA, GZMB, PRF1, and CCL3, together with several interferon-responsive genes (Figure 4C). Pathway analysis identified several gene sets uniquely enriched in active MS CD8^+^ cytotoxic T cells, including T-cell differentiation, activation, immunological synapse formation, and regulation of cytotoxic responses (Figure 4D). Tregs and PD1^+^ Tem/effector helper T cells also showed distinct transcriptional responses to EBV at Day 7, with differences in activation, proliferation, and antiviral-associated genes between active MS and HC (Supplementary Figure S8B, C). Pathways enriched across all three T-cell populations in active MS included interferon alpha/beta signaling, interferon signaling, defense response to virus, regulation of viral processes, and IL-27-mediated signaling (Figure 4E, Supplementary Figure S8D). Pathways enriched in HC relative to MS were shared across PD1^+^ Tem/effector helper T cells, Tregs, and CD8^+^ cytotoxic T cells (Supplementary Figure S8E, F).

EBV transcripts were detected across several T-cell populations, although at lower levels than in B cells (Supplementary Figure S9A, B). EBV transcript detection increased at Day 7 across several T-cell populations and subsequently declined by Day 15. Among these populations, PD1^+^ Tem/effector helper T cells showed the highest EBV positivity at Day 7, while lower levels were observed in Tregs and CD8^+^ cytotoxic T cells.

At Day15, exhaustion-associated transcriptional scores increased in selected T-cell populations from active MS donors (Figure 4F). The highest scores at Day 15 were observed in PD1^+^ Tem/effector helper T cells, Tregs, and CD8^+^ cytotoxic T cells, whereas naive CD4^+^ T cells, conventional CD8^+^ T cells, and Th17 cells showed relatively low scores. The populations with increased exhaustion-associated scores were also among those showing prominent activation and antiviral responses at Day 7. These findings indicated that T-cell responses to EBV differed between active MS and HC, with greater cytotoxic and activation-associated responses at Day 7 followed by increased exhaustion-associated transcriptional signatures at Day 15 in active MS.

### Increased EBV lytic transcription and expansion of EBV+ Age-associated B-cells in active MS

At later time points, EBV-positive cells became increasingly concentrated within the B-cell compartment, particularly at Day 15 (Supplementary Figure S3B). Given the greater accumulation of EBV-positive B cells in active MS, we further characterized B-cell populations and their transcriptional responses to EBV (Figure 5). The overall B-cell composition differed between active MS and HC, with a higher proportion of plasma cells in active MS (31% versus 14%). Age-associated B cells (ABCs) were defined using CellTypist, with reference markers including FCRL2, ITGAX, and TBX21, together with FCRL3 and FCRL4. ABCs accounted for 7% of the B-cell population in active MS but remained rare in HC (Figure 5A). Donors with stable MS showed a similar B-cell distribution, with increased representation of plasma cells and ABCs compared with HC, although both populations were less prominent than in active MS (Supplementary Figure S10A). As shown in Figure 2A, EBV positivity was also higher in ABCs, plasmablasts, and plasma cells from active MS donors compared with HC, indicating that the later EBV-associated differences in MS were most apparent within differentiated B-cell populations.

**Figure 5.**
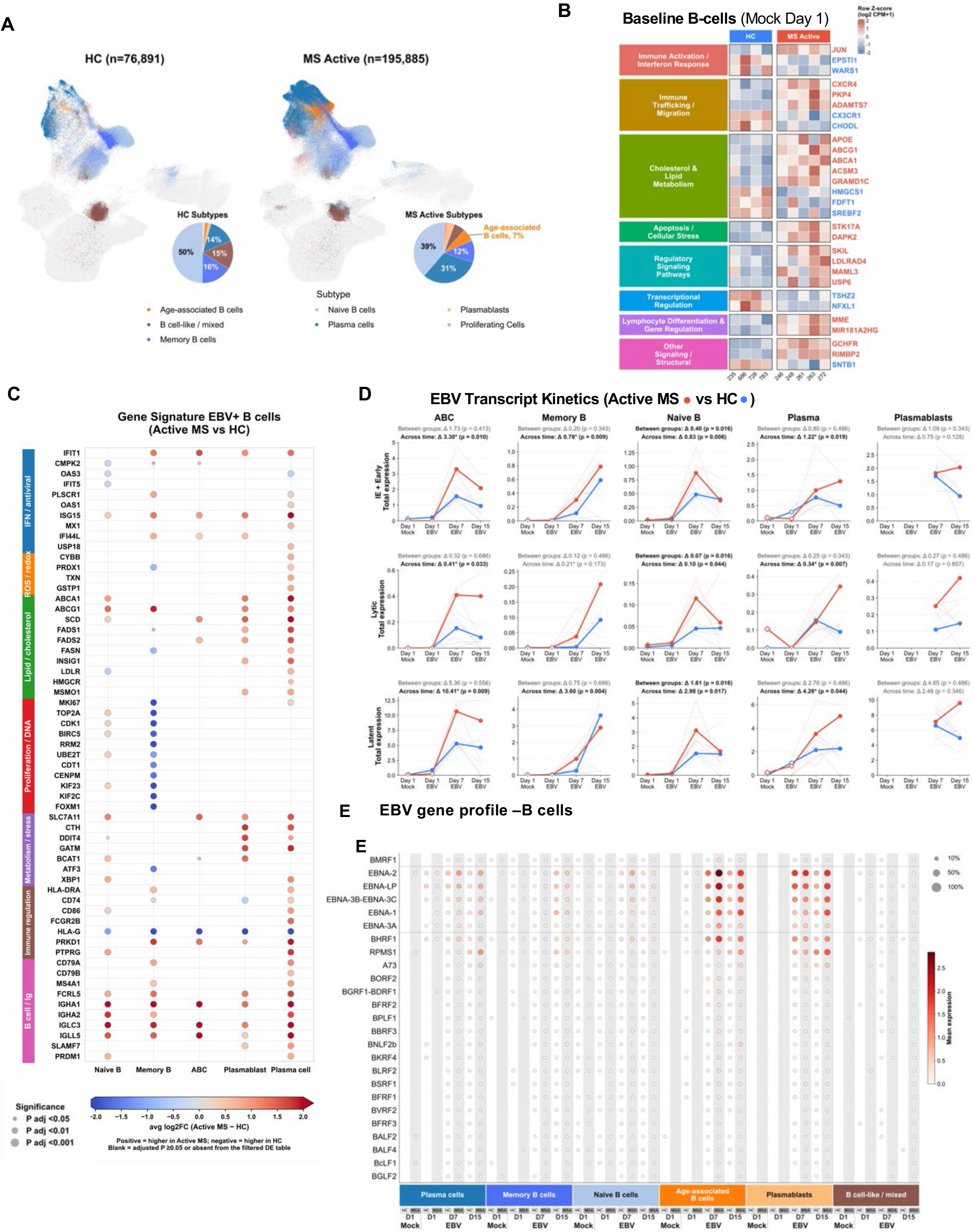
Baseline transcriptional differences and distinct EBV-associated host and viral expression profiles in B cells from active MS donors. (A) UMAP embeddings showing B-cell populations in healthy controls (HC) and active MS donors, colored by annotated subtype. Inset pie charts indicate subtype proportions. (B) Row Z-scored pseudobulk expression heatmap of genes differentially expressed between Active MS and HC B cells under baseline mock conditions on day 1. Columns represent donors, and genes are grouped by functional category. (C) Differential host gene expression between Active MS and HC within EBV-positive B-cell subtypes. Color indicates log₂ fold change (active MS versus HC), with red and blue indicating higher and lower expression in active MS, respectively. Dot size indicates statistical significance. Genes are grouped by functional category. (D) Temporal profiles of total early lytic, late lytic, and latent EBV gene expression in age-associated B cells (ABC), memory B cells, naive B cells, plasma cells, and plasmablasts. Red and blue indicate active MS and HC, respectively. Thin lines represent individual donors, and bold lines indicate group trajectories. Between-group and across-time comparisons are indicated. (E) EBV gene expression profiles across B-cell subtypes, disease groups, and conditions. Dot size indicates the percentage of cells expressing each viral gene, and color indicates mean expression.

Transcriptional differences between active MS and HC B cells were also present before EBV exposure. At baseline (Mock Day 1), B cells from active MS donors showed differences in genes associated with immune activation, cellular migration, lipid metabolism, stress responses, and transcriptional regulation (Figure 5B). Genes involved in cholesterol and lipid metabolism, including APOE, ABCG1, ABCA1, ACSM3, and GRAMD1C, were elevated in active MS, whereas HMGCS1, FDFT1, and SREBF2 showed higher expression in HC. Differences were also observed in immune trafficking genes, including CXCR4, PKP4, and ADAMTS7, together with genes associated with cellular stress and regulatory signaling.

Following EBV infection, EBV-positive B cells showed additional transcriptional differences between active MS and HC that varied across B-cell subtypes (Figure 5C). Interferon-responsive genes, including IFIT1, ISG15, MX1, and IFI44L, showed differences across memory B cells, ABCs, plasmablasts, and plasma cells. Lipid metabolism-associated genes, including ABCA1, ABCG1, SCD, FADS1, and FADS2, were also differentially expressed, with several showing higher expression in active MS. In contrast, genes associated with proliferation and cell-cycle regulation, including MKI67, TOP2A, CDK1, and CENPM, were lower in memory B cells from active MS donors. Differences in immune regulation and antigen presentation were also observed, including reduced HLA-G expression across several B-cell populations. Notably, immunoglobulin-associated genes, including IGHA1, IGHA2, IGLC3, and IGLL5, showed higher expression in active MS across multiple B-cell populations. Comparison of EBV-positive and EBV-negative cells further showed differences between active MS and HC, particularly within memory B cells, with interferon-responsive genes showing opposite directions of change between the two groups (Supplementary Figure S10B-C).

EBV transcript kinetics differed across B-cell populations and time points, with distinct patterns of early, late, and latent viral gene expression between active MS and HC (Figure 5D). Lytic-associated transcripts (e.g., BZLF1, BRLF1, BALF2, BALF5, BMRF1, and BMRF2) showed a marked increase by Day 7 in ABCs, followed by a decline at Day 15, whereas memory B cells and plasma cells showed a more gradual increase over time. Late lytic-associated transcripts (e.g., BLLF1, BcLF1, BALF4, and BFRF3) were generally detected at lower levels but showed greater abundance in active MS, particularly within ABCs, plasma cells, and plasmablasts at later time points. Latent viral transcripts (e.g., EBNA1, EBNA2, EBNA-LP, EBNA3A/B/C, LMP1, LMP2A, and LMP2B) were more abundant across B-cell populations, with a pronounced Day 7 increase in ABCs and naive B cells and more sustained expression in memory B cells and plasma cells.

EBV gene expression was further characterized across B-cell populations, with the 25 most highly expressed viral genes in B cells (Figure 5E). Latency-associated transcripts, including EBNA2, EBNA-LP, EBNA3A/B/C, EBNA1, and RPMS1, as well as BHRF1, were detected across multiple B-cell subtypes, with higher expression in active MS compared with HC. ABCs from active MS donors showed a greater viral transcriptional signal, whereas HC ABCs generally showed lower expression. Lytic-associated genes were detected less frequently and showed more variable expression across B-cell populations.

To further examine the relationship between EBV transcriptional programs and B-cell differentiation, we performed PrimeFlow RNA analysis of EBV transcripts together with surface CD27 and CD38 expression. At 72 hours following EBV exposure, cells expressing EBV early/late lytic, latency II/III, or EBER transcripts showed a greater representation of CD27+CD38+ antibody-secreting cells (ASCs) compared with transcript-negative populations, whereas immediate-early lytic transcript detection showed a smaller difference (Supplementary Figure S12). These results further supported an association between EBV transcriptional activity and antibody-secreting B-cell phenotypes following infection.

Together, these results showed that B cells from active MS donors differed from HC at baseline and showed distinct host and viral transcriptional profiles following infection. These differences were particularly apparent within differentiated B-cell populations, including plasma cells and ABCs. Given the prominent ABC response following *ex vivo* EBV infection, we further characterized ABC expansion and transcriptional differences between active MS and HC.

### EBV-positive ABCs in active MS show distinct host transcriptional states and increased lytic-associated viral expression

We further characterized ABC expansion and EBV-associated transcriptional changes in active MS. Longitudinal analysis showed a marked increase in ABC frequency following EBV infection in active MS, whereas ABC frequencies remained relatively low in HC (group × time interaction, p = 0.0015; Supplementary Figure S11C). EBV transcript abundance within ABCs also increased substantially over time in active MS (p = 0.010; Figure 6A). Given the increased EBV transcript abundance observed in ABCs from active MS donors, we further characterized lytic-associated viral gene expression within this population. Early lytic genes, including BZLF1, BRLF1, BALF2, BALF5, BMRF1, and BMRF2, and late lytic-associated genes, including BLLF1, BALF4, BcLF1, and BFRF3, were detected and showed greater expression in EBV-positive ABCs from active MS than HC (Figure 6C). The lytic module score was also higher in active MS compared with HC in EBV-positive ABCs pooled across Days 1, 7, and 15 (p = 0.040; Figure 6D).

**Figure 6.**
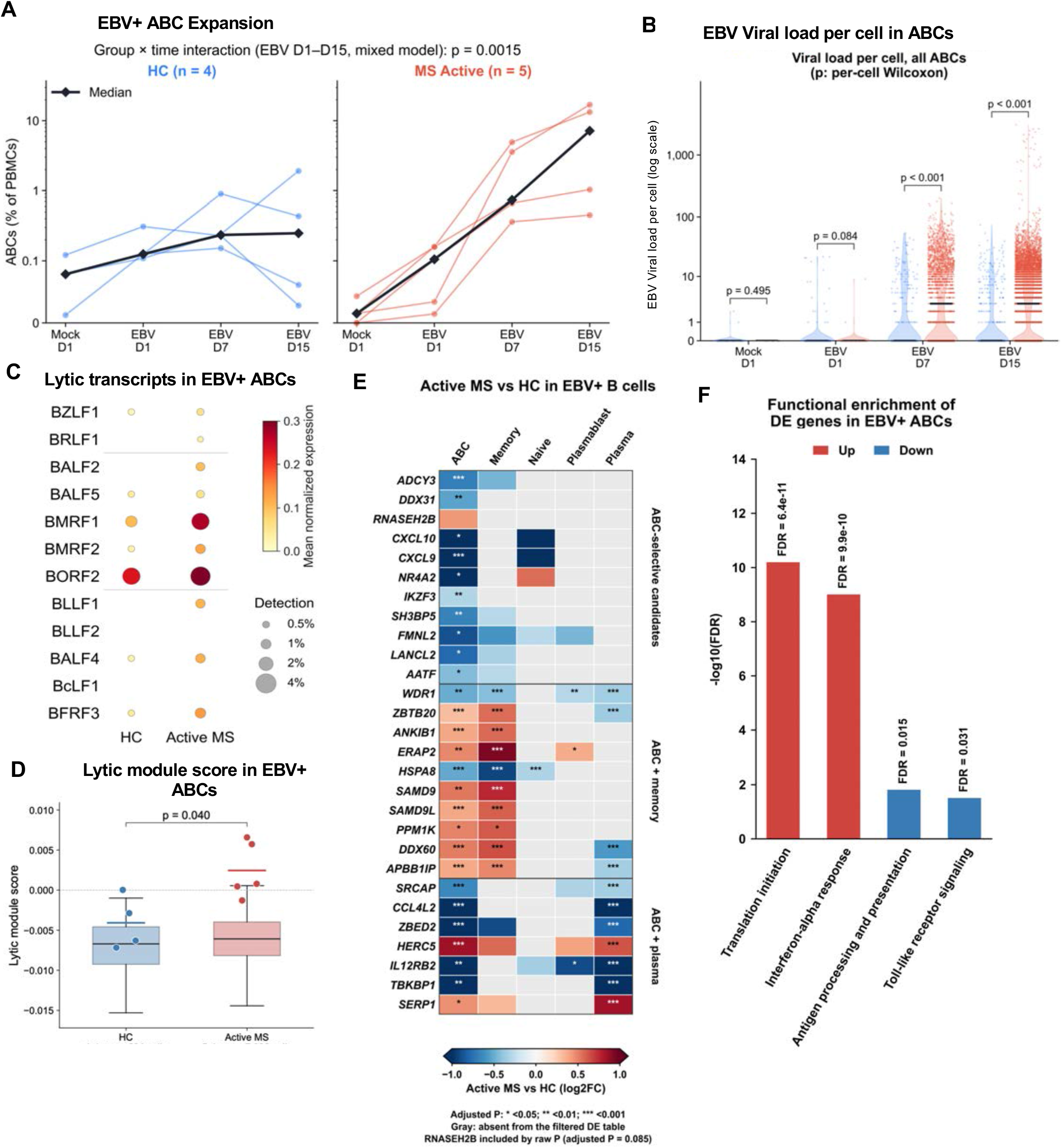
Expansion and distinct host and viral transcriptional profiles of EBV-positive age-associated B cells in active MS donors. (A) Frequencies of age-associated B cells (ABCs) among PBMCs across the indicated conditions and time points in healthy controls (HC; n = 4) and active MS donors (n = 5). Colored lines represent individual donors, and black lines indicate group medians. The group-by-time interaction across EBV-exposed samples from days 1–15 was assessed using a mixed-effects model (P = 0.0015). (B) EBV-derived UMI counts per ABC across conditions and time points. Blue and red indicate HC and Active MS, respectively. Between-group comparisons were assessed using per-cell Wilcoxon tests; P values are indicated. (C) Expression of top lytic EBV genes in EBV-positive ABCs from HC and active MS donors. Dot size indicates the percentage of cells expressing each gene, and color indicates mean normalized expression. (D) Lytic EBV module scores in EBV-positive ABCs from HC and Active MS donors. Points represent individual donors. Statistical significance was assessed using a two-sided Mann–Whitney U test, P value is indicated. (E) Differential host gene expression between active MS and HC within EBV-positive B-cell subtypes. Genes are grouped according to significant results differentially expressed in ABCs or shared with memory B cells or plasma cells. Color indicates log₂ fold change (active MS versus HC); gray indicates no result reported. Asterisks indicate adjusted P values: *P <0.05, **P <0.01, and ***P <0.001. (F) Functional enrichment of genes upregulated (red) or downregulated (blue) in EBV-positive ABCs from active MS donors compared with HC. Bars show −log₁₀(FDR); FDR-adjusted P values are indicated.

To further characterize transcriptional differences in ABCs, we examined genes identified as differentially expressed between active MS and HC within EBV-positive ABCs and compared these with other B-cell populations (Figure 6E; Supplementary Figure S11A). Several genes, including ADCY3, DDX31, CXCL9, CXCL10, IKZF3, and FMNL2, showed reduced expression in ABCs and met the differential-expression threshold only in this population. Other transcriptional differences were shared with memory B cells, including increased expression of ZBTB20, ANKIB1, ERAP2, SAMD9, and PPM1K. In contrast, ABCs shared a separate set of transcriptional changes with plasma cells, including reduced expression of CCL4L2, ZBED2, IL12RB2, and TBKBP1, together with increased expression of HERC5 and SERP1. Pathway analysis of differentially expressed genes in EBV-positive ABCs identified enrichment of translation initiation and interferon-alpha response among upregulated genes, and antigen processing and presentation and Toll-like receptor signaling among downregulated genes (Figure 6F). Overall, EBV-positive ABCs from active MS donors showed transcriptional changes detected selectively in ABCs and features shared with memory B cells and plasma cells, together with increased lytic-associated EBV transcription compared with HC.

An overview of the longitudinal gene expression patterns across immune cell compartments revealed distinct temporal immune states in response to EBV between active MS and HC (Figure 7A, Supplementary Figure S13 –14). Cellular gene trajectories across time and cell types revealed that monocyte- and NK-cell-associated transcriptional responses, particularly inflammatory and interferon-stimulated genes were highly elevated in MS compared to HC at Day 1 (Supplementary Figure S15). By Day 7, broader NK-, T-, and B-cell responses were observed in MS, followed by predominantly T- and B-cell-associated transcriptional changes at Day 15, with increasing overlap between these populations, particularly in active MS. These changes were accompanied by greater accumulation of EBV-positive B cells, particularly ABCs and plasma cells, in active MS. Taken together, these results showed that preexisting immune cell differences in MS were accompanied by distinct host and viral transcriptional responses following EBV exposure, involving early innate responses and later changes in T- and B-cell populations.

**Figure 7.**
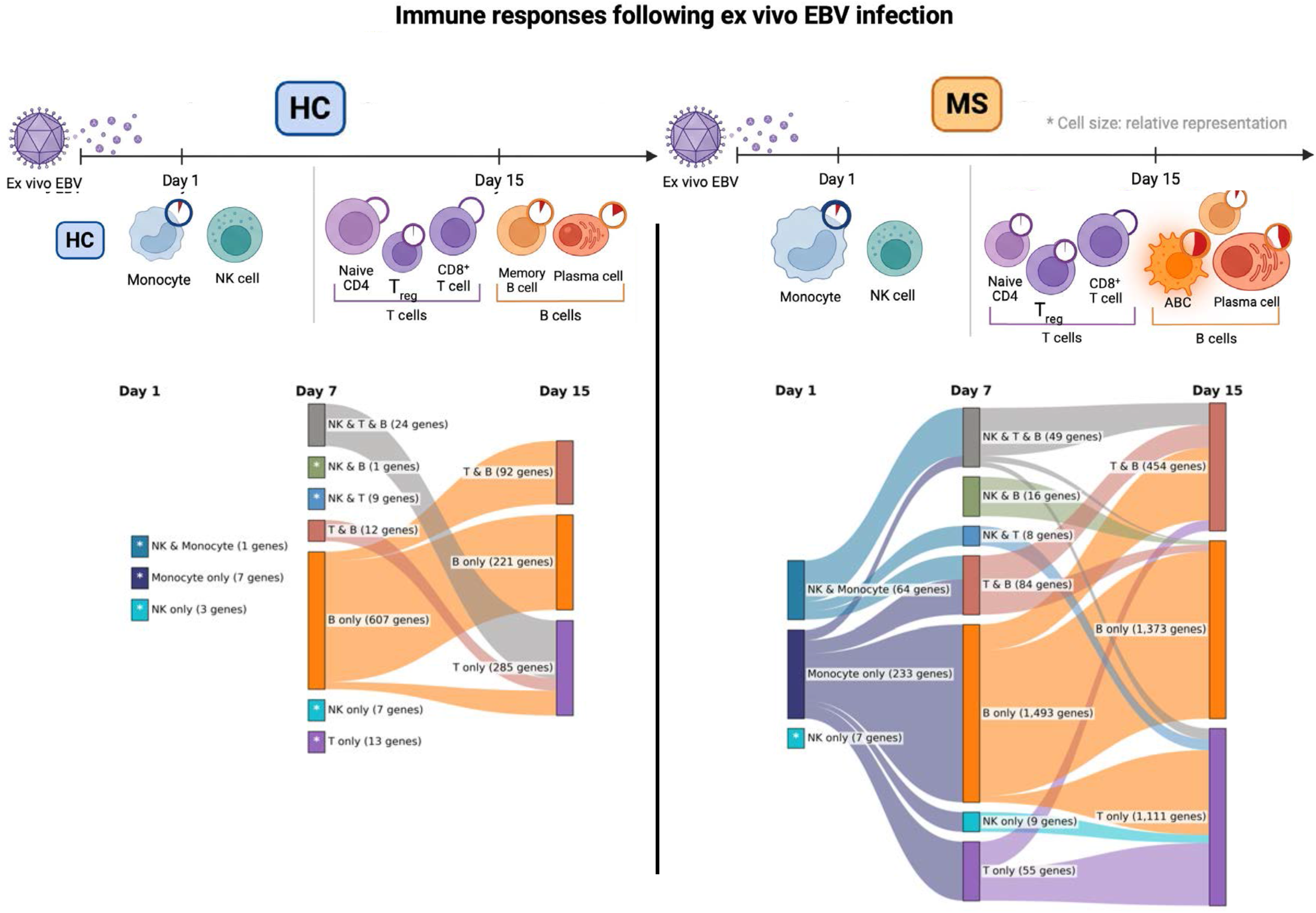
Summary of immune-cell responses following ex vivo EBV infection in healthy controls and active MS donors. Schematic illustrating early monocyte and NK-cell responses and subsequent changes in T- and B-cell populations, including increased representation of EBV-positive ABCs and plasma cells in active MS donors. Cell size indicates relative population representation, and red pie sectors indicate EBV positivity. Sankey diagrams summarizing EBV-induced gene sets across immune-cell lineages on Days 1, 7, and 15 following EBV exposure. Nodes represent lineage-specific or shared EBV-induced gene sets, with labels indicating the number of genes in each node. Connecting ribbons indicate genes retained between adjacent time points.

## DISCUSSION

In this study, we characterized the immune cell states and longitudinal responses to *ex vivo* EBV infection in PBMCs from patients with MS and HC. We identified multiple MS-associated differences, including a reduced capacity to control EBV-driven LCL outgrowth among active MS. We also detected differences in baseline immune cell states in MS that included detection of EBV transcripts in monocyte/activated macrophage subpopulations. At early time points after EBV infection, monocytes and NK cells from patients with active MS showed enhanced antiviral responses, while at later time points T-cell populations showed differences in cytotoxic and activation-associated programs, followed by increased exhaustion-associated transcriptional signatures. Within the B-cell compartment, active MS showed greater accumulation of EBV-positive cells, particularly ABCs and plasma cells, together with differences in host and viral transcriptional profiles. Notably, EBV-positive ABCs from active MS donors also showed increased lytic-associated viral gene expression compared with HC. These results suggest that preexisting differences in immune cell populations and transcriptional states may contribute to the impaired immune control of EBV and expansion of EBV-infected B-cells. This study further demonstrates the utility of the *ex vivo* PBMC infection model for investigation of patient-specific responses to EBV infection, viral-host interactions and control of EBV-driven B-cell immortalization.

Single-cell RNA sequencing provides a powerful approach for resolving EBV-associated responses across heterogeneous immune populations. Recent studies have used single-cell approaches to characterize EBV-positive B cells in autoimmune diseases. In systemic lupus erythematosus (SLE), single-cell sequencing targeting EBV identified increased frequencies of EBV-positive B cells, predominantly within CD27^+^CD21^low^ memory B cell populations, accompanied by enhanced B-cell activation, interferon signaling, and antigen-presentation programs^27^. Similar analysis of blood and CSF from patients with MS identified EBV-positive B cells with increased activation and antigen-presentation programs, including cells with ABC-like phenotypes^28^ and EBV-reactive T-cell states associated with CNS infiltration and inflammatory interactions with B-lineage and myeloid cells ^29^. Other single-cell studies have identified expansion of TBX21^+^CD21^low^CXCR3^+^ B cells associated with early MS disease activity ^20^ and EBV-associated transcriptional changes in B cells and monocytes preceding clinical relapse^23^. However, conventional single-cell approaches have limited sensitivity for detecting low-abundance EBV transcripts. To address this limitation, we incorporated targeted enrichment of EBV transcripts from the same barcoded single-cell libraries to increase viral transcript detection while maintaining the corresponding whole-transcriptome cell identity, allowing host and viral responses to be examined across different immune populations and time points. Additionally, we examined the time course of EBV infection and immune responses using a controlled *ex vivo* infection model, enabling direct comparison of MS and HC immune cell responses to a standardized EBV exposure. This approach provides a framework for identifying disease-associated differences in viral transcription and host immune responses that may influence the outcome of EBV infection.

Among the most striking differences between MS and HC were preexisting differences in monocyte and NK states that preceded *ex vivo* EBV infection. Monocytes from patients with active MS showed differences in inflammatory, lipid metabolism, and cell-migration-associated genes at baseline, followed by greater induction of interferon and antiviral pathways at Day 1, particularly within macrophage-like monocyte populations. EBV transcripts were also detected in activated/phagocytic and macrophage-like monocytes at early time points, with greater EBV positivity and increased frequencies of CD14^+^gp350^+^ cells in MS compared with HC. Previous studies have shown that EBV can induce ROS production ^30,31^ and promote inflammatory and immunosuppressive responses in monocyte-derived macrophages through TNF-α, IL-6, and IDO signaling^32^. More recently, mitochondrial ROS produced by monocyte-derived macrophages was shown to contribute directly to neurological tissue damage during neuroinflammation^33^. Although EBV transcript detection in monocytes does not establish productive infection and may instead reflect viral uptake, phagocytosis, or processing of infected cellular material, the preferential detection of viral transcripts within activated monocyte subsets suggests that viral association is influenced by cellular states. EBV infection of monocytes has been reported previously, including in the context of HIV-positive oral hairy leukoplakia characterized by poorly controlled EBV lytic infection, as well as during *ex vivo* infection of purified monocyte-derived macrophages^32,34^. The greater EBV positivity within specific activated monocyte populations may also suggest that EBV infection promotes inflammatory monocyte activation. This would be consistent with the known role of myeloid cells in B-cell activation, oxidative stress, and neuroinflammation in MS^35^.

NK cells also showed differences in baseline composition, with reduced CD16^+^ NK-cell frequencies in active MS despite a stronger antiviral transcriptional response at Day 1, potentially reflecting a distinction between cytokine-mediated antiviral activation and direct cytotoxic immune surveillance. NK cells can respond to inflammatory cytokines, including type I interferons, IL-12, and IL-18, without directly recognizing infected cells^36^. In addition, previous work has shown that EBV-associated immune evasion through HLA-E can limit NK-cell-mediated elimination of autoreactive cells in MS^37,38^. Thus, the stronger interferon-associated transcriptional response observed in active MS may not necessarily translate into more effective elimination of EBV-infected B cells. By Day 7, both active MS and HC showed induction of overlapping antiviral genes and pathways, consistent with an earlier antiviral transcriptional response in active MS. Interestingly, EBV transcripts were detected predominantly in CD16^−^ NK cells, whereas CD16^+^ NK cells showed strong antiviral gene induction without detectable viral RNA. This difference may reflect indirect activation through interferon signaling or interactions with EBV-exposed immune cells, rather than direct viral association. Such cytokine-driven responses may contribute to early antiviral activation without necessarily enhancing the cytotoxic elimination of EBV-infected B cells, although this remains to be determined.

Differences in T-cell responses emerged more prominently at later time points. At Day 7, CD8^+^ cytotoxic T cells from active MS donors showed greater induction of cytotoxic and activation-associated genes, including GZMA, GZMB, PRF1, and CCL3, despite broadly overlapping antiviral transcriptional responses between MS and HC populations. Previous studies have identified expanded T-cell populations recognizing EBV-infected B cells in the CSF of patients with MS^39^, while more recent work has demonstrated clonally expanded EBV-reactive CD8^+^ T cells in MS^40^, supporting a role for EBV-directed cytotoxic T-cell responses in MS. HLA-B7-restricted EBV-specific CD8^+^ T cells with exhaustion-associated phenotypes have also been reported^41^. In our study, exhaustion-associated transcriptional scores increased at Day 15 in several T-cell populations from active MS donors, particularly PD1^+^ Tem/effector helper T cells, Tregs, and CD8^+^ cytotoxic T cells. The greater cytotoxic response at Day 7 and increased exhaustion-associated transcriptional signatures at Day 15 may reflect sustained activation followed by increased inhibitory regulation during prolonged EBV exposure. In Tregs, however, increased exhaustion-associated scores may also reflect activation or a suppressive transcriptional state rather than functional exhaustion. These responses could potentially limit the effectiveness of EBV-specific immune surveillance and contribute to increased accumulation of EBV-positive B cells in MS. However, because exhaustion-associated transcriptional signatures do not necessarily establish functional T-cell exhaustion, additional studies examining functional T-cell antigen specificity, cytotoxic function, and inhibitory receptor signaling will be needed to determine whether these changes reflect impaired antiviral immunity.

The coexistence of enhanced antiviral and cytotoxic transcriptional responses with greater accumulation of EBV-positive B cells in active MS suggests that heightened immune activation does not necessarily correspond to effective control of EBV-infected cells. Exploratory analysis of matched bulk RNA-seq and clinical data further identified inverse associations between early EBV-induced B-cell differentiation-associated responses and CSF IgG index, which was not observed under baseline Mock conditions. Together with the baseline transcriptional differences observed across immune populations, these findings raise the possibility that pre-existing immune states influence peripheral responses to EBV and may be associated with intrathecal humoral immune activity and subsequent clinical outcomes. However, these exploratory associations require further validation, and additional mechanisms involving altered B-cell states or sustained immune regulation may contribute to EBV-associated B-cell expansion in MS.

The most pronounced differences at later time points occurred within the B-cell compartment, where active MS patients exhibited greater accumulation of EBV-positive cells, particularly ABCs, plasmablasts, and plasma cells. The expansion of the ABC population is especially relevant given their proposed role in MS-associated immune dysregulation. Previous studies have identified expanded T-bet^+^CXCR3^+^ atypical B cell populations in MS that may contribute to inflammatory B-/T-cell interactions and CNS immune cell recruitment^19,21^. In our study, EBV-positive ABCs showed distinct transcriptional differences between active MS and HC, including changes detected selectively in ABCs and changes shared with memory B cells and plasma cells. These included genes involved in antigen processing (ERAP2), interferon-associated ISGylation (HERC5), and inflammatory regulation (CXCL9, and CXCL10). These differences raise the possibility that ABCs in active MS have altered capacities for antigen presentation and immune signaling, and CNS-trafficking, which may influence their interactions with T cells and the surrounding immune environment. Comparisons of EBV-positive and EBV-negative cells further showed that EBV-associated transcriptional changes differed across B-cell subtypes, involving interferon responses, proliferation, lipid metabolism, and differentiation-associated programs. These findings suggest that B-cell responses to EBV in active MS are influenced by distinct cellular states, with altered immune signaling and differentiation-associated transcriptional programs in ABCs and plasma cells.

EBV transcript patterns in active MS reflected differences in B-cell differentiation states. Increased detection of EBNA2 and other latency-associated transcripts in differentiated B cells from active MS donors may be relevant to the altered host transcriptional profiles observed in these populations. EBNA2 can regulate B-cell activation and survival through interactions with host transcription factors and MS-associated genetic susceptibility loci^24,42^. Conversely, plasma-cell differentiation can promote EBV lytic reactivation through activation of the immediate-early viral genes BZLF1 and BRLF1, involving differentiation-associated transcription factors such as BLIMP1^15^. Interestingly, EBV-positive ABCs from active MS donors showed increased viral transcript abundance and lytic-associated gene expression compared with HC, consistent with previous reports of increased EBV lytic transcription in B cells derived from patients with active MS^25^. Among the viral lytic genes upregulated in MS were BMRF1 (viral DNA processivity factor) and BORF2 (viral ribonucleotide reductase and inhibitor of APOBEC3B)^43,44^, raising the possibility that altered viral DNA replication and APOBEC-associated mutagenic processes may influence EBV-infected B-cell states. These observations suggest that altered B-cell differentiation and viral transcriptional regulation may be linked in active MS, with ABCs showing both prominent latency-associated gene detection and increased lytic-associated transcription.

Several limitations should be considered. Although longitudinal *ex vivo* EBV infection allowed us to compare immune responses under controlled experimental conditions, this model does not fully reproduce the complex interactions involved in persistent EBV infection *in vivo*. Although differences in immune cell composition were observed under Mock conditions, these measurements were obtained following PBMC thawing and culture and may not fully reflect the immune cell distributions in peripheral blood prior to culture. In addition, detection of EBV transcripts within individual immune populations does not necessarily indicate productive infection, particularly in non-B-cell populations. Differences in immune cell composition and transcriptional responses may also be influenced by donor-specific factors, including disease activity. Further studies will be needed to determine whether the altered antiviral responses and increased latent and lytic-associated transcription observed in our PBMC model contribute directly to ongoing inflammation in MS.

Collectively, our findings demonstrate the utility of an *ex vivo* PBMC infection model for investigating dysregulated immune responses to EBV in MS. We identified distinct baseline immune cell states and longitudinal responses to EBV exposure in patients with MS compared with healthy controls, revealing disease-associated transcriptional signatures across both innate and adaptive immune populations. Enhanced early antiviral responses and increased detection of EBV transcripts within activated monocyte populations suggest that interactions between EBV and non-B-cell populations may contribute to immune dysregulation in MS. We further observed preferential expansion of EBV-positive ABCs, accompanied by distinct host transcriptional programs and increased lytic-associated viral gene expression, providing new insight into the potential contribution of these cells to MS pathogenesis. These changes occurred in conjunction with heightened T-cell activation followed by exhaustion-associated transcriptional signatures, indicating that increased immune activation does not necessarily translate into effective control of EBV-infected cells. Together, these findings support a model in which altered innate and adaptive immune responses to EBV promote the expansion of pathogenic B-cell populations and sustained inflammatory programs that may contribute to disease activity and progression in MS. Future studies leveraging this *ex vivo* system to prevent the emergence of EBV-positive ABCs or attenuate EBV-associated inflammatory responses may further define mechanisms linking EBV-driven immune dysregulation to MS pathogenesis and facilitate the development of therapies targeting EBV-associated disease processes.

## METHODS

### PBMC sample collection

Blood samples were collected from patients and healthy controls at the University of Pennsylvania Multiple Sclerosis and Related Disorders Clinic. This study was reviewed and approved by the Institutional Review Board (IRB) / Ethics Committees of the University of Pennsylvania (protocol approval number: 816805) and The Wistar Institute (protocol approval number: 22307360). Written informed consent was voluntarily obtained from all participating human subjects prior to their enrollment in the study and any study-related procedures. Participants were fully informed of the study’s purpose, the procedures involved, including the risks and discomforts associated with venous blood draw (phlebotomy), and their right to withdraw from the study at any time without penalty or loss of medical care. Further, to protect participant confidentiality, all collected blood samples and derived Peripheral Blood Mononuclear Cells (PBMCs) were anonymized and assigned a unique alphanumeric code. Data and samples were stored securely in password-protected databases and restricted-access liquid nitrogen freezers, accessible only to authorized study personnel.

Venous blood samples (120 mL per donor) were collected from patients with multiple sclerosis (MS) and healthy controls at the University of Pennsylvania in 10 mL K2EDTA tubes and processed within 4 hours of collection. Peripheral blood mononuclear cells (PBMCs) were isolated by density-gradient centrifugation using Ficoll (GE Healthcare) or Lymphoprep (STEMCELL Technologies). Whole blood was diluted 1:1 with PBS, layered over 15 mL of density-gradient medium, and centrifuged at 1,200 × g for 10 minutes at room temperature without brake. The mononuclear cell fraction was collected and washed with PBS before cryopreservation in heat-inactivated human AB serum supplemented with 10% DMSO or CryoStor CS10 (BioLife Solutions), depending on the sample preparation protocol.

Patients with MS were classified as active MS (AMS) based on the presence of at least one gadolinium-enhancing lesion on brain or spinal cord MRI within three months prior to blood collection, whereas stable MS (SMS) patients had no gadolinium-enhancing lesions during this period. Disease activity was classified according to MRI findings, as previously described^25^, with the sampling interval adapted to the current cohort.

### Clinical data collection

Clinical information, including disease activity, MRI findings, treatment history, and Expanded Disability Status Scale (EDSS) scores, was obtained from available clinical records. Laboratory measurements, including anti-EBNA1 antibody levels, CSF IgG index, neurofilament light chain (NfL), and EBV DNA levels, were obtained from available clinical and laboratory records. Clinical measurements were matched to individual donors using their study identifiers.

### EBV infection

For *ex vivo* EBV infection, cryopreserved PBMCs were thawed in RPMI 1640 supplemented with 10% FBS, 1% GlutaMAX, 100 U/mL penicillin, and 100 μg/mL streptomycin. Cells were allowed to recover overnight at 37°C in a humidified incubator with 5% CO₂ and subsequently plated in 12-well plates at a density of 1 × 10 cells/mL. PBMCs were infected with the type I EBV strain Mutu I at a multiplicity of infection (MOI) of 1 or maintained under matched mock conditions. Samples were collected at Days 1 and 7 for bulk RNA sequencing and at Days 1, 7, and 15 for single-cell RNA sequencing. PBMCs used for bulk and single-cell RNA sequencing were thawed and processed on separate occasions.

EBV stocks were generated from Mutu I cells following induction of the lytic cycle with sodium butyrate (2 μM) and 12-O-tetradecanoylphorbol-13-acetate (TPA; 20 ng/mL) for 72 hours, as previously described^26^. Virus-containing supernatants were clarified, filtered through a 0.45 μm membrane, and concentrated by ultracentrifugation through a 10% sucrose cushion at 27,000 × g for 90 minutes at 4°C. Viral DNA was quantified by droplet digital PCR (ddPCR) targeting EBV LMP1 using the QX200 system (Bio-Rad), following the previously established protocol^25^.

### Flow cytometry

Flow cytometry was performed to characterize monocyte populations and assess EBV gp350 detection following ex vivo infection. PBMCs collected under Mock and EBV-infected conditions were stained with Zombie-NIR (BioLegend) or LIVE/DEAD Fixable Aqua (Thermo Fisher Scientific) for 20 minutes at room temperature, followed by incubation with Human TruStain FcX Fc receptor blocking reagent (BioLegend) for 20 minutes on ice. Cells were subsequently stained with fluorochrome-conjugated antibodies in FACS buffer (PBS containing 5% FBS and 0.1% sodium azide) for 30 minutes at 4°C.

The monocyte panel included antibodies against CD14 (BUV395, clone MφP9, BD Biosciences), CD16 (BV711, clone B73.1, BioLegend), CD11c (BUV661, clone B-ly6, BD Biosciences), CD64 (APC, clone S18012C, BioLegend), HLA-DR (PE-Cy7, clone L243, BioLegend), CCR1 (PE, clone 5F10B29, BioLegend), CCR2 (APC-Fire, clone K036C2, BioLegend), and EBV gp350 (FITC, clone C61H, Invitrogen). Following staining, cells were washed twice with FACS buffer and resuspended for acquisition. Samples were acquired using a BD FACSymphony A3 flow cytometer (BD Biosciences). Compensation was performed using single-stained controls and compensation beads. Data were analyzed using FlowJo software (v10, BD Biosciences). Doublets and nonviable cells were excluded before identification of monocyte populations. Monocytes were identified based on CD14, CD16, and HLA-DR expression, and EBV-associated cells were assessed by gp350 staining. CCR1- and CCR2-expressing monocyte populations were quantified within the corresponding gated populations. Gating strategies are shown in Supplementary Figure S6.

### Bulk RNA sequencing

Total RNA was extracted from Mock- and EBV-infected PBMCs collected at the indicated time points using the RNeasy Mini Kit (Qiagen, Cat# 74104), including on-column DNase treatment according to the manufacturer’s protocol. RNA quality was assessed using an Agilent TapeStation with all samples showing RNA integrity numbers (RIN) greater than 7. Sequencing libraries were prepared using the QuantSeq 3′ mRNA-Seq Library Prep Kit (Lexogen) according to the manufacturer’s instructions. Libraries were sequenced on an Illumina NovaSeq X platform. Sequencing reads were aligned to a combined reference containing the human genome (hg19) and EBV genome (NC_007605.1) using STAR ^45^ ^45^. Gene-level read counts and reads per kilobase per million mapped reads (RPKM) were estimated using RSEM (v1.2.12) ^46^with Ensembl gene annotations (GRCh37.p13). Differential gene expression analysis was performed using DESeq2 ^47^ based on raw read counts. Normalized expression values were used for downstream visualization, and differentially expressed genes were analyzed for functional enrichment using Enrichr^48^. Principal component analysis, heatmaps, and other visualizations were generated in R using DESeq2, pheatmap, and ggplot2.

### Bulk RNA-seq signature scoring and clinical correlation analysis

Raw gene counts were normalized using the median-ratio size-factor method with the positive-count estimator (poscounts), followed by log₂ transformation with a pseudocount of 1. For each donor, EBV-induced transcriptional responses were calculated as the difference in log₂(normalized counts + 1) between matched EBV-exposed and Mock samples at the corresponding time point. B-cell/antibody-secreting cell (ASC)-associated and interferon-associated transcriptional scores were calculated as the mean expression difference across selected genes. The B-cell/ASC signature included PRDM1, XBP1, IRF4, MZB1, JCHAIN, IGHA1, IGHG1, SDC1, CD38, and TNFRSF17. Donor-level scores were correlated with CSF IgG index score using two-sided Spearman rank correlation tests.

### Single-cell RNA sequencing

Single-cell RNA sequencing libraries were prepared using the Evercode Whole Transcriptome Mega Kit (Parse Biosciences, Seattle, WA, USA), based on split-pool combinatorial barcoding technology. PBMCs from patients with active MS, stable MS, and healthy controls were collected at Days 1, 7, and 15 following ex vivo EBV infection, with matched Mock controls collected at Days 1 and 7. Cells were stained with LIVE/DEAD Fixable Aqua viability dye (Thermo Fisher Scientific), and viable cells were isolated by fluorescence-activated cell sorting (FACS) to exclude dead cells prior to fixation. Approximately 12,000–20,000 cells were targeted per sample. Cells were fixed and processed according to the manufacturer’s protocol. Fixed cells underwent three rounds of split-pool combinatorial barcoding. During the first round, each sample was assigned to separate wells of a 96-well plate, allowing the first-round barcode to identify the sample of origin. Cells from all samples were subsequently pooled and randomly redistributed for the second and third rounds of barcoding, generating unique barcode combinations for individual cells. Following barcoding, cells were lysed, and whole-transcriptome libraries were prepared, amplified, and purified according to the manufacturer’s instructions. Library quality and concentration were assessed prior to sequencing. Whole-transcriptome libraries were sequenced on an Illumina NovaSeq X platform, targeting a minimum sequencing depth of 20,000 reads per cell, with approximately 25 billion reads generated in total. The same barcoded libraries were subsequently used for targeted EBV transcript enrichment.

### Targeted EBV transcript enrichment by hybridization capture

To increase the sensitivity of EBV transcript detection at single-cell resolution, targeted hybridization capture was performed using custom-designed biotinylated probes spanning the EBV genome (NC_007605.1). Whole-transcriptome libraries generated using the Parse Biosciences Evercode WT Mega Kit were pooled and subjected to hybridization capture to enrich EBV-derived sequences. Following hybridization, probe-bound library fragments were isolated using streptavidin-coated magnetic beads, washed to remove nonspecifically bound fragments, and amplified by PCR according to the capture protocol. Enriched libraries were purified and assessed for library size and concentration prior to sequencing. EBV-enriched libraries were sequenced on an Illumina NovaSeq X platform using 150-bp paired-end reads (2 × 150 bp), targeting a minimum sequencing depth of 20,000 read pairs per cell. Because targeted enrichment was performed on the same barcoded libraries used for whole-transcriptome sequencing, the original combinatorial cell barcodes were retained. EBV transcripts detected in the enriched libraries were subsequently assigned to their corresponding cells in the whole-transcriptome dataset, allowing viral gene expression to be analyzed together with host transcriptional profiles and cell-type annotations. Because targeted EBV enrichment involved hybridization capture followed by streptavidin pull-down and washing, not all WT cell barcodes were recovered in the enriched libraries. In addition, recovery of a cell barcode in the enriched library did not necessarily indicate EBV positivity due to nonspecific capture and background signals.

### Data analysis Library alignment

Reads were aligned and quantified with split-pipe against a combined reference of GRCh38 (GENCODE release 45) and the EBV type 1 genome (NC_007605.1; 88 viral genes and transcripts). The 17 sequencing libraries were processed in two split-pipe combine runs, recorded for each cell as its processing batch.

### Quality control and preprocessing

WT cells with fewer than 200 detected genes or fewer than 500 UMIs were removed, leaving 1,287,503 cells (median 1,627 genes and 2,581 UMIs per cell). Enriched-library cells were retained with at least 50 detected genes (256,067 cells), a lower threshold because targeted enrichment libraries detect far fewer features per cell.

Expression was normalized to 10,000 UMIs per cell and log-transformed. PCA on the 2,000 most variable genes, a 30-neighbor graph on 30 components, UMAP ^49^ ^50^ and Leiden clustering^51^ were computed on the WT data. Enriched cells were projected onto the WT embedding with scanpy.tl.ingest.

### Count matrices for differential expression

The analysis object stored log-normalized expression, x = log(1 + 10,000 × c / T), and each cell’s total UMI count T. Integer host-gene counts were reconstructed for DESeq2 as c = round(expm1(x) × T / 10,000). The reconstruction was exact for all 1,287,503 WT cells: every value lay within 0.007 of an integer (median per-cell maximum deviation 1.5 × 10⁻), and each cell’s counts summed exactly to its stored UMI total. Viral counts were taken directly as UMI counts from the enriched-library.

### Linking WT and enriched cells

The two split-pipe combine runs assign barcodes independently, so the same sample-and-barcode label can occur in both. Cells were therefore identified by a three-part key: sample, processing batch and the full split-pool barcode, including the sub-library index. This key was unique for every WT and every enriched cell, and 99.99% of enriched cells (256,031) linked one-to-one to a WT cell. Without the batch term, 33,858 spurious cross-run matches would have arisen. The pairing of processing batches between libraries was confirmed from the matched cells.

Cell identity always came from the WT library and viral measurements from the enriched library. Host-gene analyses used WT cells only.

### Viral load and EBV-positive cells

Per-gene viral UMI counts were taken from the raw enriched-library alignments. EBNA-LP (repeated W exons in IR1) and LMP2A (unique first exon; remaining exons shared with LMP2B) were counted at the exon level. A cell was EBV-positive when it carried at least 2 EBV UMIs from at least 2 distinct EBV genes; this excludes signal confined to a single viral gene, which can arise from ambient RNA or sequencing artifacts. By this definition, 71,910 of 256,067 enriched cells (28.1%) were EBV-positive.

Each WT cell inherited the viral counts and EBV status of its linked enriched cell. Viral readouts were total viral load (EBV UMIs per group and time point), the percentage of EBV-positive cells per donor, and EBV gene expression, summarized either over EBV-positive cells ("profile") or over all WT cells of a type, with unlinked or uninfected cells counted as zero ("burden").

### EBV gene classes

- **Immediate-early/early lytic:** BZLF1, BRLF1, BALF2, BALF5, BBLF1, BBLF4, BGLF4, BGLF5, BHRF1, BMRF1, BMRF2, BORF2, BSLF1, BaRF1.
- **Late lytic:** BLLF1, BLLF2, BcLF1, BdRF1, BFRF3, BXRF1, BVRF2, BALF4, BILF2.
- **Latent:** EBNA1, EBNA2, EBNA-LP, EBNA3A, EBNA3B/3C, LMP1, LMP2A, LMP2B, RPMS1, A73.

EBNA3B and EBNA3C share a single feature in the reference and were counted once.

### Cell-type annotation

Cell types were first predicted with CellTypist (Immune_All_Low model, majority voting) and reviewed against canonical markers. Tissue-type predictions such as macrophages were merged into the corresponding monocyte states, cytotoxic effector-memory T-cell subsets were merged into one CD8^+^ cytotoxic T-cell state, and cycling cells were labelled "proliferating cells". Two clusters with monocyte profiles but T-cell labels were reassigned to monocyte states.

NK cells, CD8^+^ cytotoxic T cells and cycling T cells, which overlapped in the embedding, were re-clustered separately and reassigned by marker expression. The Tcm/naive helper T-cell population, the largest in the dataset, was sub-typed by high-resolution clustering of a representative subset. Clusters were assigned by lineage markers (CD8A/CD8B, CD4, MS4A1/CD79A, LYZ/S100A9, FOXP3) and by naive (CD4, IL7R, CCR7, TCF7) versus memory (CD4, IL7R, S100A4, CD40LG) marker scores, giving naive CD4+, memory CD4+, CD8+, regulatory T and mixed states. Labels were transferred to the remaining cells by nearest-neighbor voting.

The final annotation has 29 cell states. Mixed or atypical states, which probably include doublets, were kept as separate categories and not interpreted as populations. Enriched cells took the label of their linked WT cell.

### Pseudobulk differential expression

Raw counts of WT cells were summed per donor, condition and cell type (decoupler), excluding EBV genes^52^. Donors contributing fewer than 50 cells to a contrast (20 for B-cell subtypes) were excluded, and each contrast required at least two donors per side. Genes with fewer than 10 total counts were removed.

DESeq2 ^47,53^ was fitted with median-of-ratios size factors, empirical-Bayes dispersion shrinkage and Cook’s-distance refitting. Contrasts were tested by the Wald test with Benjamini–Hochberg adjustment; log2 fold changes are unshrunk. Genes were called differentially expressed at adjusted p < 0.05 and |log2 fold change| > 0.5, which defines the genes highlighted in volcano plots and those used for pathway analysis. A stricter |log2 fold change| ≥ 1 was used for EBV-induced genes in the gene cascade, for DEG counts in the B-cell and ABC summary tables, and, with adjusted p ≤ 0.01, for gene lists comparing T-cell responses between groups. One HC donor’s day-15 EBV sample was excluded from EBV-response analyses because its culture formed an LCL (EBV positivity 48.8% vs 3.8-14.8% in other HC donors).

### Contrasts

1. **Baseline disease effect:** MS vs HC in mock day-1 cultures. The three main monocyte states were also analyzed together.
2. **EBV response within each disease group:** EBV vs mock at days 1, 7 and 15, fitted separately in HC and MS. Side-by-side volcano and pathway plots show these separate fits, not a between-group test.
3. **Disease effect within a condition:** MS vs HC within mock and infected cultures at each time point. This was the main design for six T-cell subtypes (PD-1^+^ effector helper T, regulatory T, naive CD4^+^ T, CD8^+^ cytotoxic T, CD8^+^ T and type 17 helper T cells).
4. **Difference in EBV response between groups:** interaction model ∼group + infection + group:infection. In T cells, interaction tests found few significant genes, as expected with four to five donors per group. T-cell group differences were therefore assessed by design 3 and by the difference between the groups’ EBV-induced log2 fold changes from design 2 ("delta-delta"), a descriptive effect size without its own test.
5. **EBV-positive vs EBV-negative cells within infected donors** (B cells): paired design ∼donor + status.

### Pathway analysis

Up- and down-regulated genes of each contrast (at least 10 genes) were tested separately for over-representation. Up to 500 genes, ranked by adjusted p, were submitted to Enrichr ^48,54,55^ through GSEApy and tested against GO Biological Process 2023, Reactome 2022 and KEGG 2021. B-cell and ABC analyses used local MSigDB Hallmark and GO Biological Process libraries, with all genes tested in the contrast as background. Terms with adjusted p < 0.05 were retained.

Plots show each contrast’s top terms by adjusted p, with Enrichr combined score ^55^ or −log10 adjusted p as bar length. Where HC and active MS EBV responses are shown side by side, each group’s terms were ranked independently, and the plot shows the union of both top lists.

Pre-ranked GSEA ^56^ used each contrast’s Wald statistic (GSEApy; 1,000 permutations; gene-set size 15–500), with significance at FDR q < 0.05.

### EBV-induced gene cascade

For each disease group, EBV-vs-mock DESeq2 was fitted per cell subtype and time point (a node): NK and monocyte subtypes at day 1, and T- and B-cell subtypes at days 7 and 15.

Monocytes contributed day-1 nodes only, because few remained in later cultures; plasma cells and plasmablasts were not analyzed, because they are essentially absent from mock cultures. A gene was induced at a node at adjusted p < 0.05 and log2FC ≥ 1.

Each cascade gene was assigned a broad functional category by majority vote over its GO Biological Process annotations (GOATOOLS), ignoring annotations that describe only lineage differentiation or development. Six interferon-stimulated genes lacking an interferon- or virus-related annotation (CMPK2, LY6E, HELZ2, NRIR, SPATS2L, SAMD9L) were assigned to the antiviral/interferon category by curation.

### Gene-signature scoring and cell composition

Signature scores were computed per cell type with scanpy.tl.score_genes against expression-matched control genes (control set ≥ 50 genes)^57^. The T-cell exhaustion signature comprised TOX, CTLA4, PDCD1, HAVCR2, LAG3 and TIGIT. Donor means used donor–cell type–condition groups with at least 50 cells.

EBV lytic module scores were calculated using Seurat’s AddModuleScore based on 12 EBV lytic genes (BZLF1, BRLF1, BALF2, BALF5, BMRF1, BMRF2, BORF2, BLLF1, BLLF2, BALF4, BcLF1, and BFRF3). Normalized expression of the selected genes was averaged and compared with expression-matched control genes selected from 24 expression bins, with up to 100 controls per gene. Scores were averaged per donor across Days 1, 7, and 15, and differences between active MS and HC were assessed using a two-sided exact permutation test on donor-level means.

Composition was computed from WT cells as the percentage of each donor’s cells per condition, requiring at least 20 cells. For the composition figure comparing HC with MS, active and stable MS donors were combined.

### Visualization

UMAPs show WT cells with lineage density contours. Time courses are plotted at equal spacing; main figures compare HC with active MS, and supplementary figures include stable MS. Percentages spanning several orders of magnitude, including zero, are plotted on symmetric-log axes, which are linear near zero. A cell state is shown separately when it reached at least 200 cells or 1% of cells in a displayed condition; hollow markers indicate values from fewer than 200 cells. Trend-line panels show the cell states with the largest between-group difference or within-group change. In summary heatmaps, lighter cells indicate values from a single donor.

### Quantification and statistical analysis

Donors were the unit of replication; n (donors) is given in each figure legend, and analyses with cells as observations are stated explicitly. Tests were two-sided, and p < 0.05 (or adjusted p / FDR q < 0.05 for multiple tests) was considered significant. Multiple testing was controlled by the Benjamini–Hochberg procedure within each DE contrast and pathway library.

Donor-level values at a single time point were compared by exact Mann–Whitney U (two groups) or Kruskal–Wallis^58^ (three groups). Signature scores were compared by one-way ANOVA, with Tukey’s HSD for three groups and Cohen’s d as effect size. EBV positivity and viral load were compared per time point by one-way ANOVA on donor-level values; pooled positivity rates of two groups within a cell type and time point were compared by Fisher’s exact test^59^.

ABC frequencies (% of PBMCs) over EBV days 1, 7 and 15 were log10-transformed (pseudocount 0.01 percentage points) and modelled with a linear mixed model, y ∼ day × group + (1 | donor) (statsmodels, REML). The group × day interaction was tested by Wald and likelihood-ratio tests. Per-donor expansion rates (slope of log10 ABC % per day, shown as fold change per week) were compared by Welch’s t-test, an exact permutation test and Mann–Whitney U.

ABC viral load was compared per donor by Mann–Whitney U and a mixed model on log10(total + 1), and per cell by Wilcoxon rank-sum tests (cells as observations) complemented by a mixed model with a donor random intercept.

### PrimeFlow: Single cell surface immunophenotyping combined with fluorescence in situ hybridization (FISH) of EBV lytic and latent transcripts

To determine whether EBV drives B cells into antibody-secreting cell populations, we used Primeflow during a short-term infection timecourse (Mutu virus, MOI=1 at 0, 12, 24, 48, and 72h) of thawed PBMC (1 × 10 PBMCs) from healthy donors. Cells were stained with viability dye and an optimized antibody panel targeting B cell and lineage-specific markers (live-dead, aqua; CD19 BUV496; IgD, BV480; CD27 BV650; and CD38 BUV 395). RNA-FISH was then performed using fluorophore-conjugated probes specific for EBV transcripts (Table Y). EBV+ and EBV–B cells were immunophenotyped and categorized into canonical subsets based on surface marker expression. Assay performance was validated using EBV+ and EBV-cell lines and fluorescence minus one control. Data were acquired on an Aurora (Cytek), spectrally unmixed using single-stained reference controls, and analyzed using OMIQ software (Dotmatics).

**Table Y.**
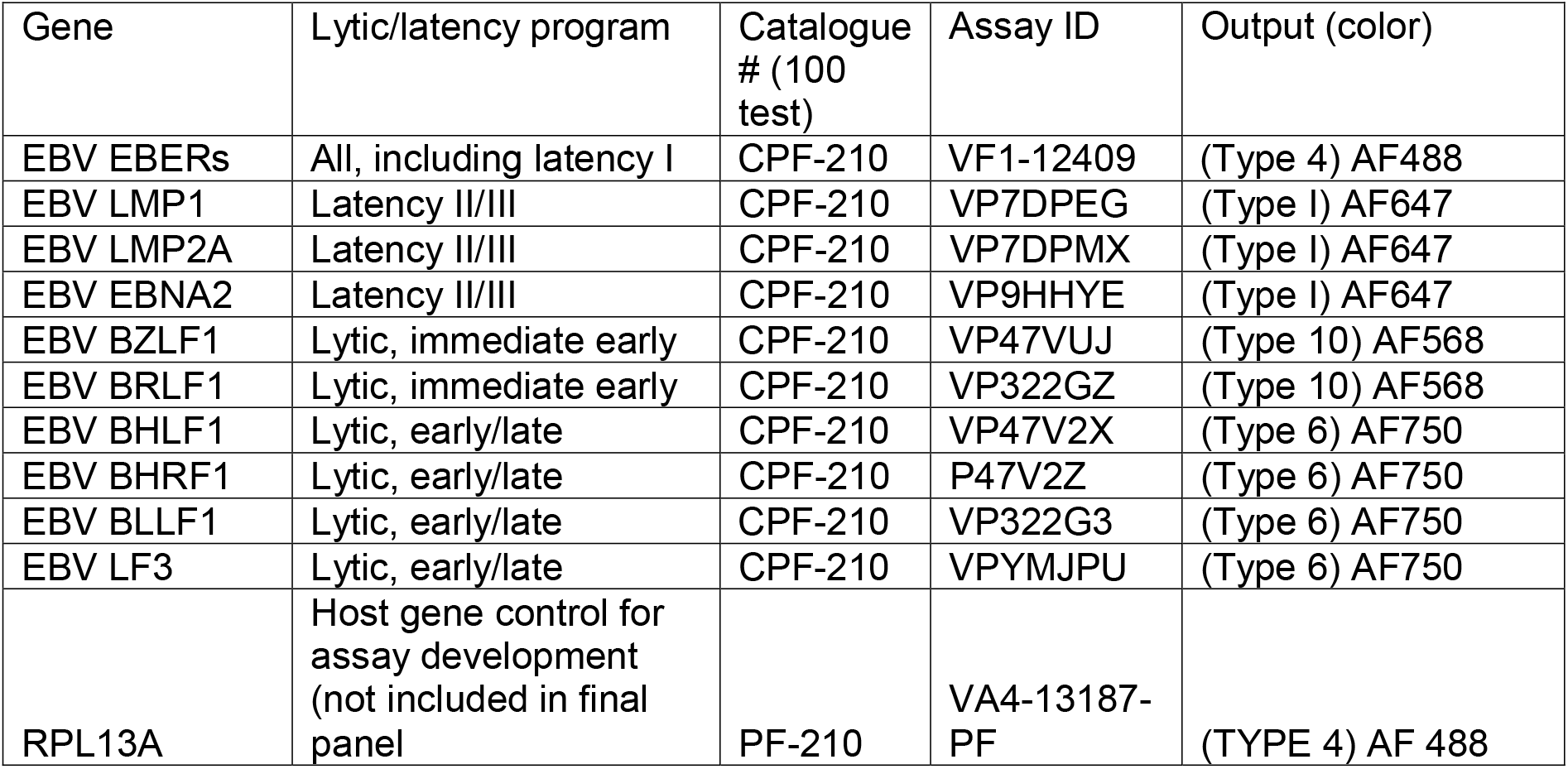
PrimeFLow Probes.

## Data Availability

RNA-sequencing data generated in this study have been deposited in the Gene Expression Omnibus (GEO) under accession GSEXXXX. Source data underlying the graphical representations in the main and Extended Data figures are provided with the paper.

## Code availability

No custom software was developed for this study. Statistical and computational analyses were performed using standard software and open-source packages as described in the Methods.

## Ethics and Inclusion statement

This study was approved by the Institutional Review Boards of the University of Pennsylvania (Protocol 816805). Written informed consent was obtained from all participants in accordance with the Declaration of Helsinki and institutional requirements. No bias or discriminatory language was used in the study design, data analysis, or manuscript writing. All contributors have been appropriately acknowledged, and authorship reflects substantial contribution in accordance with the ICMJE criteria.

## ACKNOWLEDGEMENTS

We thank members of the Lieberman and Bar-Or laboratories for their assistance throughout. This work was supported by grants from the NIH R01 AI153508, R01 DE017336, P01 CA281867 to PML, United States Department of Defense Award Number: HT9425-23-1-1049 (SSS and PML), U01 CA284811 (RFA and RRX).

## Competing Interests

The authors declare no competing interests.

## Author Contributions

## FIGURE LEGENDS

**Supplementary Table S1.** Clinical characteristics and experimental measurements of study donors. Donors are listed by Penn-PNI identifier and grouped according to clinical diagnosis, including active MS, stable MS, other MS, non-MS neurological controls, and healthy controls (HC). Experimental assays and lymphoblastoid cell line (LCL) formation outcomes following ex vivo EBV infection are indicated for each donor. Available clinical information includes demographic characteristics, EBV serology and viral measurements, cerebrospinal fluid (CSF) measurements, MRI findings, treatment history, and Expanded Disability Status Scale (EDSS) scores. Missing values indicate unavailable clinical or laboratory measurements.

## Supplementary figures

**Supplementary Figure S1**. Bulk RNA-sequencing analysis of multiple sclerosis and control cohorts. (A) Study cohorts for bulk and single-cell RNA sequencing. The bulk RNA-sequencing cohort comprised healthy controls (n = 5), individuals with multiple sclerosis (MS; n = 12), and non-MS neurological controls (n = 5). (B) Principal component analysis (PCA) of bulk RNA-seq samples across the indicated conditions and time points. Colors indicate MS and non-MS groups, and shapes indicate Days 1 and 7. (C) Hierarchically clustered heatmap of selected differentially expressed genes between MS and non-MS PBMCs under EBV-exposed culture conditions. Columns represent individual samples, and rows represent genes. Colors indicate relative gene expression, with red and blue indicating higher and lower expression, respectively. The annotation bar indicates donor group. (D) Gene Ontology (GO) Biological Process enrichment analysis using the GO Biological Process 2025 gene set collection. The x-axis indicates the combined enrichment score, dot size represents −log10(P value), and dot color indicates gene overlap. (E) Correlation between CSF IgG index and Day 1 EBV-induced B-cell/antibody-secreting cell (ASC)-associated transcriptional scores in MS donors (n = 12). (F) Correlation between CSF IgG index and Day 1 Mock B-cell/ASC-associated transcriptional scores in MS donors (n = 12). In E and F, each point represents an individual donor, and transcriptional responses were calculated as the mean difference in log₂(normalized counts + 1) between matched EBV-exposed and Mock samples. Spearman correlation coefficients and unadjusted P values are indicated. Neither association remained significant after Benjamini–Hochberg correction for multiple comparisons.

**Supplementary Figure S2.** Validation of CellTypist-based immune-cell annotations using canonical marker gene expression. Initial cell-type annotations were assigned using CellTypist and subsequently validated and refined based on canonical marker gene expression. Dot plot showing the expression of selected marker genes across annotated immune-cell populations in the combined single-cell RNA-seq dataset. Dot size represents the percentage of cells expressing each gene, and color indicates the mean normalized expression among expressing cells. Cell-type frequencies are indicated in parentheses, and marker genes are grouped by immune-cell lineage, including B cells, T cells, myeloid cells, and NK cells.

**Supplementary Figure S3.** Distribution of EBV-positive cells across immune cell populations (A) UMAP projections of whole-transcriptome (WT) PBMC data from the stable MS group under mock conditions on day 1 and following EBV exposure on days 1, 7, and 15. Cells are colored by annotated cell type, with EBV-positive cells overlaid in dark red. Inset pie charts indicate the cell-type composition of EBV-positive cells in each condition. (B) UMAP projections showing EBV-positive cells colored by annotated cell type in healthy controls (HC), stable MS, and active MS groups across the indicated conditions and time points. EBV-positive cells were defined as cells with ≥2 EBV-derived UMIs at least two distinct EBV genes. Cells not meeting these criteria are shown in gray.

**Supplementary Figure S4**. EBV transcript abundance and numbers of EBV-positive cells across disease groups and time points. (A) Total EBV-derived UMI counts normalized per 10,000 cells in healthy controls (HC) and active MS samples under mock conditions on day 1 and following EBV exposure on days 1, 7, and 15. Bars show the mean across donors and error bars the standard error of the mean (SEM); points represent individual donors (n = 4-5 per group). The y-axis is displayed on a log10 scale; means, SEM and statistical tests were calculated on the untransformed values. P values are shown above the comparisons. Solid brackets: two-sided unpaired Student’s t-test on donor values. The dotted bracket (EBV Day 15) indicates a one-sided Mann–Whitney U test comparing active MS donors with HC donors whose cultures did not generate LCLs (active MS > HC). HC donor 728 (diamond) was excluded from this comparison because its culture successfully generated an LCL. (B) Absolute numbers of EBV-positive cells by annotated cell type in HC, active MS, and stable MS groups across the indicated conditions and time points. Stacked bars show the contributions of the top 10 cell types overall.

**Supplementary Figure S5.** Monocyte populations, EBV-induced transcriptional responses, and viral gene expression. (A) UMAP projection showing monocyte and related myeloid populations in the stable MS donors, colored by annotated subtype. The inset pie chart indicates the relative proportions of these populations. (B) Differential gene expression in activated/phagocytic monocytes following EBV exposure versus mock treatment on day 1, analyzed separately within HC and active MS groups. HC results are displayed on the left and active MS results on the right. The y-axis indicates −log₁₀(adjusted P value). These represent within-group responses to EBV exposure rather than a direct comparison between active MS and HC. (C) Dot plot showing EBV gene expression profiles among EBV-positive monocytes across the indicated subtypes, disease groups, and conditions. Dot size indicates the proportion of cells expressing each gene, and color indicates average expression.

**Supplementary Figure S6.** Flow cytometric analysis of monocyte gp350 expression and chemokine receptor profiles on day 1. (A) Representative gating strategy showing selection of cells by forward and side scatter, singlet discrimination, identification of monocyte populations using CD14 and CD16, and assessment of CCR1 and CCR2 expression. (B) Representative flow cytometry plot showing gp350 and CD14 expression, colored by HLA-DR intensity. (C–E) Frequencies of CCR1⁺CCR2⁺ monocytes (C), total CCR1⁺ monocytes (D), and total CCR2⁺ monocytes (E) in mock- and EBV-exposed samples from healthy controls (HC; n = 7) and individuals with MS (n = 8) on day 1. Points represent individual donors, and lines connect paired samples. Statistical significance was assessed using paired Wilcoxon signed-rank tests; P values are indicated.

**Supplementary Figure S7.** NK-cell populations, transcriptional responses to EBV exposure, and viral gene expression. (A) UMAP showing NK-cell and related populations in the stable MS, colored by annotated subtype. The inset pie chart indicates subtype proportions. (B) Heatmaps showing the mean percentages of NK cells, CD16⁺ NK cells, and CD16⁻ NK cells among PBMCs in healthy controls (HC), active MS, and stable MS groups across the indicated conditions and time points. (C) Pathway enrichment analysis of differentially expressed genes between MS and HC CD16⁺ NK cells under baseline mock conditions on day 1. Bar length indicates the combined enrichment score, and color indicates −log₁₀(adjusted P value). (D) Differential gene expression following EBV exposure versus mock treatment on day 1 in CD16⁻ NK cells, analyzed separately within HC and active MS. (E) Overlap of EBV-induced pathways in MS among total NK cells, CD16⁺ NK cells, and CD16⁻ NK cells. (F, G) Differential gene expression following EBV exposure versus mock treatment on day 7 in CD16⁻ (F) and CD16⁺ (G) NK cells. In D, F, and G, HC results are displayed on the left and active MS results on the right; the y-axis indicates −log₁₀(adjusted P value). (H) EBV gene expression profiles in EBV-positive NK cells across the indicated groups and conditions. Dot size indicates the proportion of EBV-positive cells expressing each gene, and color indicates mean expression. (I, J) Top 10 enriched pathways associated with EBV-induced genes on day 7 in CD16⁻ (I) and CD16⁺ (J) NK cells, shown separately for HC and active MS groups. Bar length indicates −log₁₀(adjusted P value).

**Supplementary Figure S8**. T-cell populations and transcriptional responses to EBV exposure. (A) UMAP showing T-cell populations in the stable MS donors, colored by annotated subtype. The inset pie chart indicates subtype proportions. (B, C) Differential gene expression following EBV exposure versus mock treatment on day 7 in regulatory T cells (B) and Tcm/effector helper T PD1⁺ cells (C), analyzed separately within healthy controls (HC) and active MS samples. HC results are displayed on the left and active MS results on the right. The y-axis indicates −log₁₀(adjusted P value). (D, E) Overlap of pathways enriched among EBV-upregulated genes in Tcm/effector helper T PD1⁺ cells, regulatory T cells, and CD8⁺ cytotoxic T cells in active MS (D) and HC (E) on day 7. (F) HC pathways shared across all three T-cell populations, ranked by mean combined enrichment score across cell types. Color indicates −log₁₀(adjusted P value). (G, H) Pathways unique to individual T-cell populations in HC (G) and active MS (H) on day 7, restricted to terms with a combined enrichment score >500.

**Supplementary Figure S9.** EBV gene expression and positivity across T-cell populations. (A) Dot plot showing EBV gene expression profiles in EBV-positive T cells across the indicated subtypes, disease groups, and conditions. Dot size indicates the proportion of EBV-positive cells expressing each gene, and color indicates mean expression. (B) Heatmap showing the percentage of EBV-positive cells within each T-cell subtype in healthy controls (HC) and active MS samples under mock conditions on day 1 and following EBV exposure on days 1, 7, and 15. EBV-positive cells were defined as cells with ≥2 EBV-derived UMIs spanning at least two distinct EBV genes. Gray tiles indicate groups containing fewer than 20 whole-transcriptome (WT) cells and are excluded from the color scale. Percentages are displayed within tiles. Asterisks denote Fisher’s exact test comparisons between active MS and HC within the same timepoint (* p<0.05, ** p<0.01, *** p<0.001).

**Supplementary Figure S10.** B-cell populations and host transcriptional differences associated with EBV positivity. (A) UMAP of B-cell populations in the stable MS donors, colored by annotated subtype and remaining cells shown in gray. The inset pie chart indicates the subtype composition of B cell subtype proportions. (B) Differences between active MS and healthy controls (HC) in EBV-associated host gene expression across B-cell subtypes. Color represents the difference in log₂ fold change for EBV-positive versus EBV-negative cells between groups (active MS minus HC). Symbol shape indicates the direction of the EBV-positive versus EBV-negative comparison within each group. Symbols are shown for genes with FDR <0.05 in both groups. Genes are grouped by functional category. (C) Selected host gene expression comparisons across age-associated B cells (ABC), memory B cells, naive B cells, plasmablasts, and plasma cells. The left column shows log₂ fold changes between active MS and HC within EBV-positive cells. The middle column shows log₂ fold changes between EBV-positive and EBV-negative cells within each disease group. The right column shows correlations between host gene expression and EBV transcript abundance. Orange and blue points indicate active MS and HC, respectively; open circles indicate adjusted P ≥0.05. Blank entries indicate unavailable results.

**Supplementary Figure S11.** Transcriptional differences in EBV-positive age-associated B cells and related B-cell populations. (A) Heatmap showing log₂ fold changes between active MS and healthy controls (HC) within EBV-positive naive B cells, memory B cells, age-associated B cells (ABC), plasmablasts, and plasma cells. Genes are grouped by shared changes, opposite directions, or significant entries restricted to ABC or selected combinations of B-cell subtypes. Gray indicates absence from the supplied significant-results table. (B) Heatmap showing log₂ fold changes between EBV-positive and EBV-negative cells within ABC, memory B-cell, and plasma-cell populations, analyzed separately in HC and active MS groups. Selected genes are organized by descriptive functional categories. Gray indicates unavailable results. Adjusted P values: *P <0.05, **P <0.01, and ***P <0.001.

**Supplementary Figure S12.** EBV transcriptional states in antibody-secreting B cells following infection. (A) Representative contour plots showing CD27 and CD38 expression in live CD19⁺ B cells positive (red) or negative (blue) for the indicated EBV transcripts, measured by PrimeFlow RNA analysis 72 h after infection of PBMCs with Mutu-1 virus. Probe sets detect immediate-early lytic, early/late lytic, latency II/III, and EBER transcripts. Antibody-secreting cells (ASCs) were defined as CD27⁺CD38⁺ B cells. (B) Proportions of ASC⁺ (red) and ASC⁻ (blue) cells within each EBV transcript-positive or transcript-negative B-cell population at 72 h post-infection. Horizontal dotted and dashed lines indicate total ASC frequencies at baseline (0 h; 1.54%) and 72 h post-infection (41.96%), respectively.

**Supplementary Figure S13.** Shared and cell-type-specific transcriptional responses to EBV exposure. (A) Matrix showing the numbers of EBV-induced genes within and shared between the indicated cell populations and time points in healthy controls (HC) and active MS samples. Diagonal entries indicate the total numbers of induced genes in each population (active MS, orange; HC, blue). The upper-right and lower-left triangles show the numbers of shared induced genes in active MS and HC, respectively. EBV-induced genes were defined by adjusted P <0.05 and log₂ fold change ≥1 for EBV exposure versus mock treatment. (B) Donor-level pseudobulk differential expression analysis using DESeq2 in classical, macrophage-like, and activated/phagocytic monocytes on day 1, comparing EBV exposure with mock treatment separately in HC (top) and active MS (bottom). Red and blue points indicate significantly upregulated and downregulated genes, respectively (adjusted P <0.05; |log₂ fold change| ≥1). Orange points indicate significant genes with |log₂ fold change| <1, and gray points indicate nonsignificant genes. Numbers of induced genes are indicated in each plot. Y-axis scales differ between rows.

**Supplementary Figure S14.** Donor-level validation and functional classification of EBV-induced transcriptional responses. (A) Donor-level pseudobulk differential expression analysis using DESeq2, comparing EBV exposure with mock treatment on day 1 in CD16⁺ NK cells, CD16⁻ NK cells, total NK cells, macrophage-like monocytes, and activated/phagocytic monocytes. Results are shown separately for healthy controls (HC; top) and active MS samples (bottom). Red points indicate induced genes (adjusted P <0.05; log₂ fold change ≥1), blue points indicate downregulated genes (adjusted P <0.05; log₂ fold change ≤−1), orange points indicate significant genes with |log₂ fold change| <1, and gray points indicate nonsignificant genes. Numbers of induced genes are indicated in each plot. (B) Functional category composition of EBV-induced gene sets restricted to or shared among the indicated immune-cell lineages on days 1, 7, and 15, shown separately for HC and active MS. Dot size indicates gene count, and color indicates the percentage assigned to each functional category. (C) Functional category breakdown of genes detected as EBV-induced only in HC, shared between HC and active MS, or only in active MS, stratified by lineage-sharing pattern and time point. Tile colors and numbers indicate gene counts.

**Supplementary Figure S15.** Temporal distribution of EBV-positive immune cells and an MS-associated EBV-response gene signature. (A) Bubble-pie plots showing selected immune-cell populations in healthy controls (HC) and active MS samples on days 1, 7, and 15 following EBV exposure. Bubble size represents each population’s percentage within its disease group and time point. Red pie sectors indicate the percentage of EBV-positive cells within each population; the remaining sectors indicate EBV-negative cells. Ring colors denote monocyte, T-cell, and B-cell lineages. (B) Heatmap showing differences in EBV-induced gene expression between active MS and HC across the indicated cell populations (rows) and time points. EBV versus mock log_2_ fold changes (log_2_FC) were computed by pseudobulk DESeq2, separately for each group. Of the 57 genes induced by EBV (padj < 0.05, log_2_FC >= 1) in at least half of the active MS cell populations, the 14 shown are those whose log_2_FC in active MS exceeded that in HC by >= 2 in at least one population. Values represent the difference in log_2_FC (EBV versus mock; active MS minus HC). Red and blue indicate larger and smaller responses in active MS, respectively. Black outlines mark genes significantly induced in active MS but not in HC in that population. White cells indicate genes do not test in one of the groups. Column annotations classify genes as antiviral/interferon-associated, other immune, metabolic, or other.

## Notes

### Competing Interest Statement

The authors have declared no competing interest.

