## supplementary figures for "A Dysregulated Peripheral Immune Landscape Favors the Expansion of Epstein-Barr Virus-positive Age-Associated B Cells in Multiple Sclerosis"

### Supplementary Figure S1

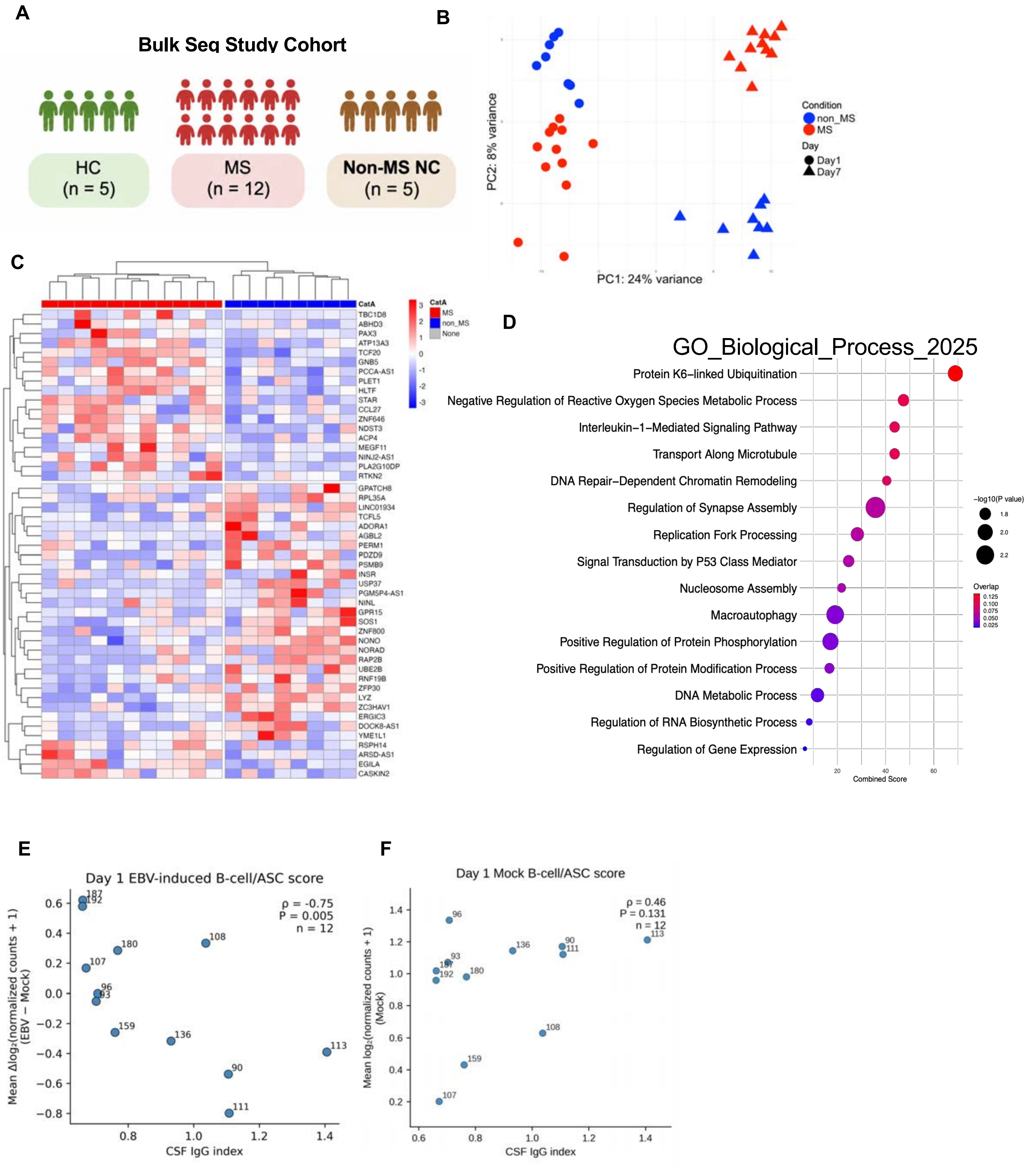

### Supplementary Figure S2

#### Cell type annotation – validation with canonical marker genes

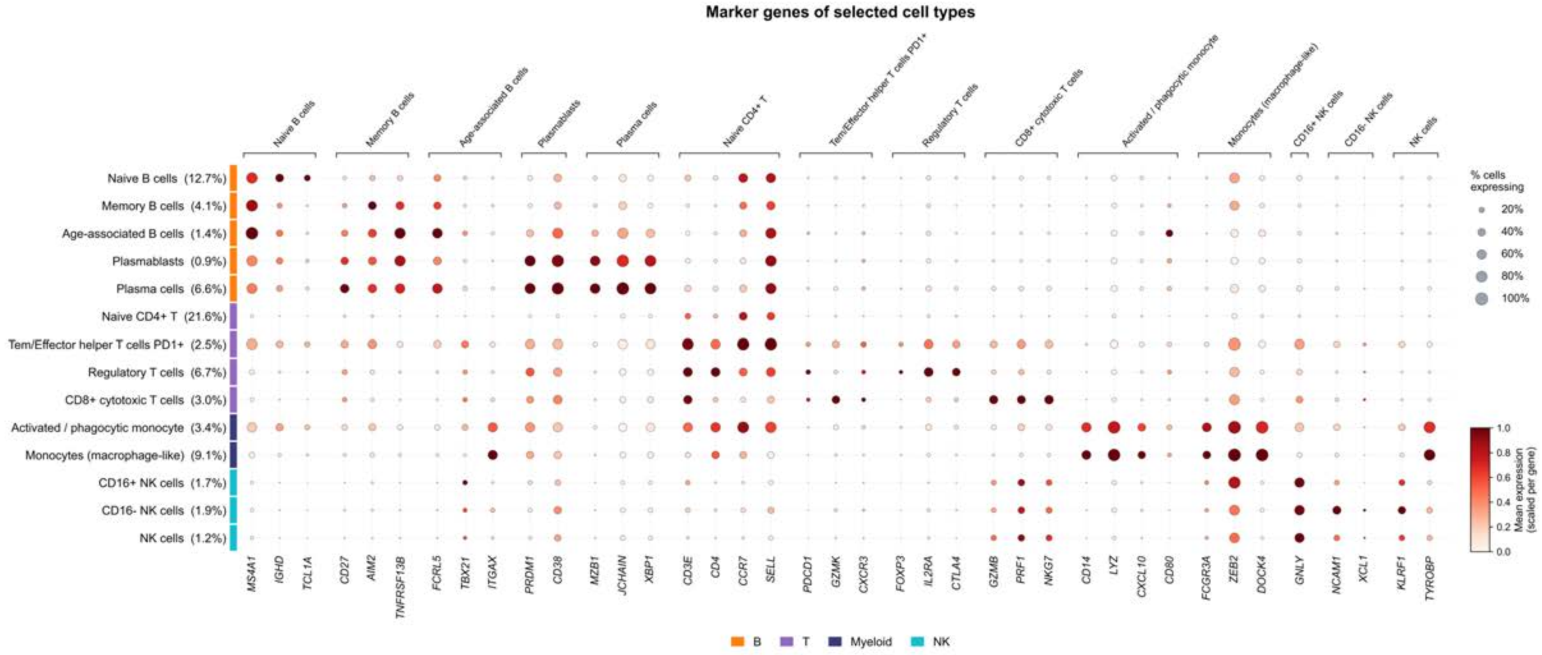

### Supplementary Figure S3

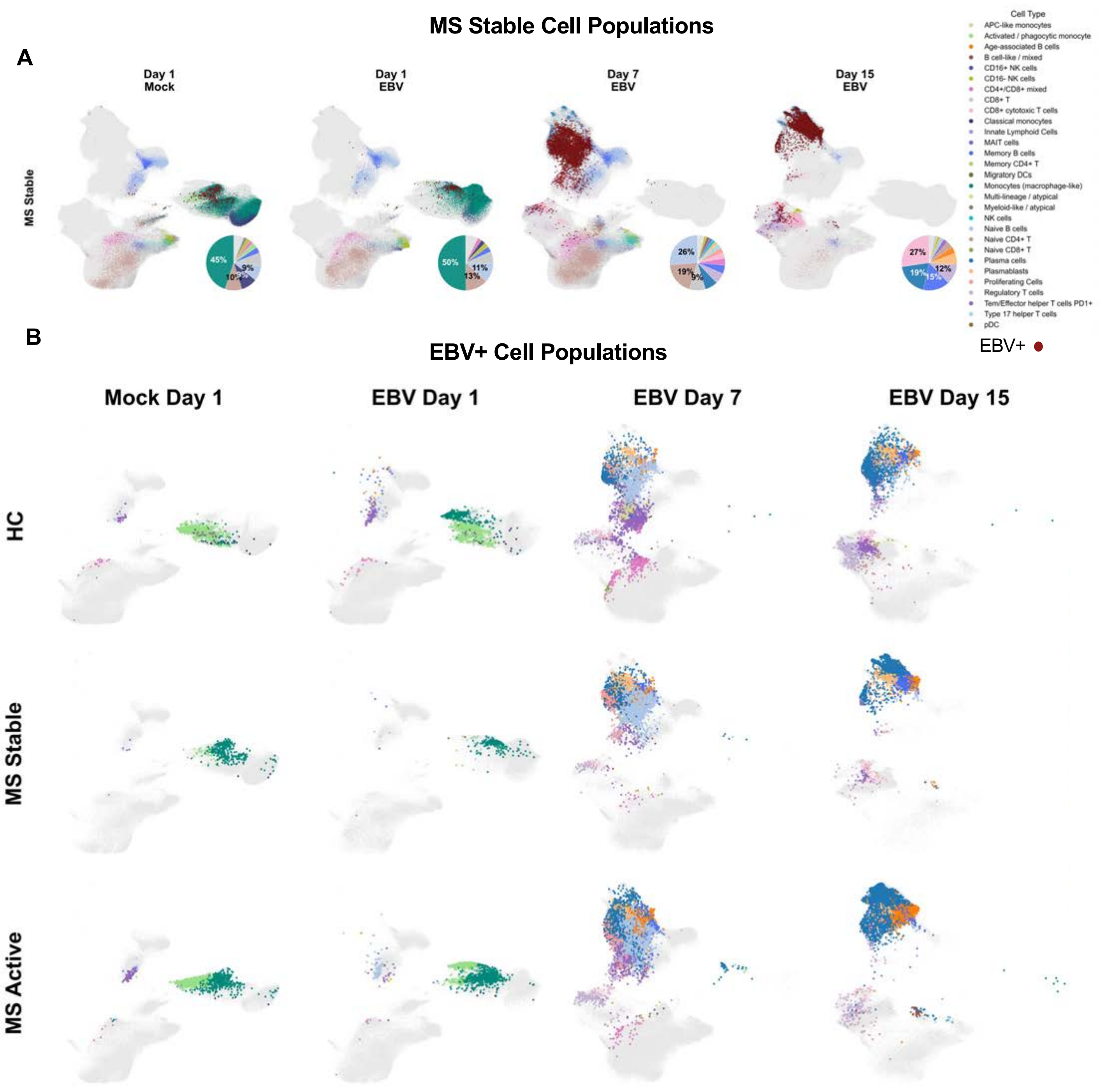

### Supplementary Figure S4

A

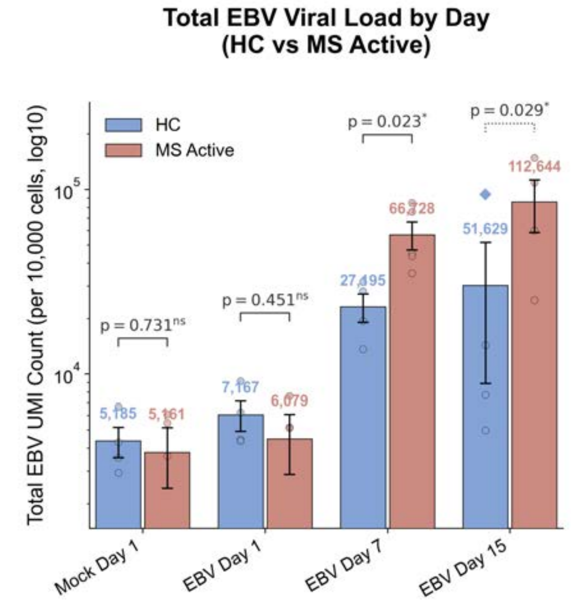


B

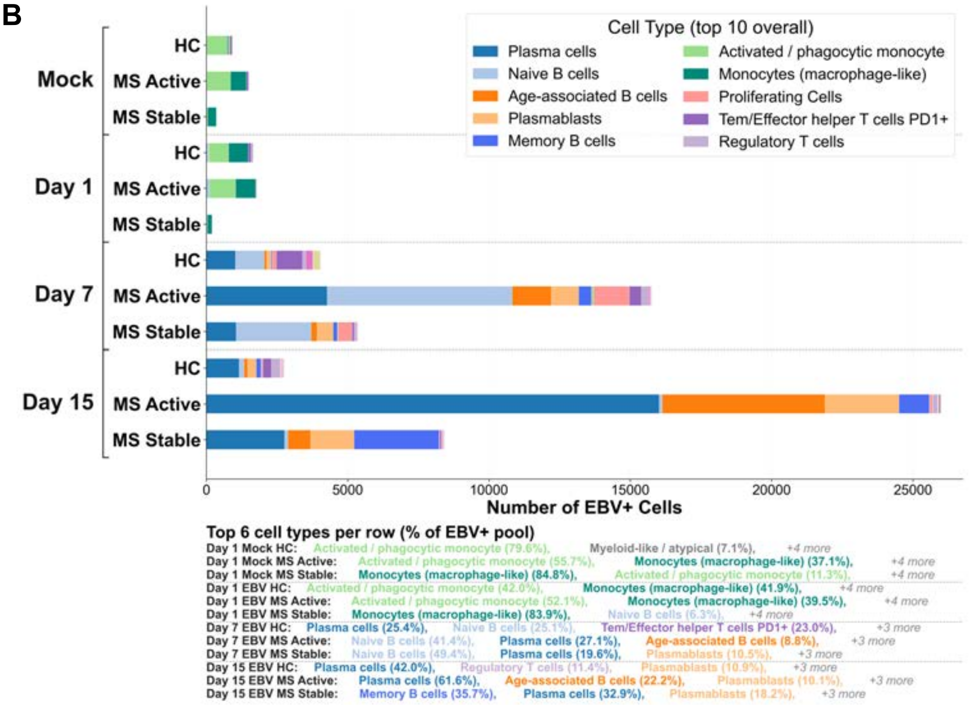

Supplementary Figure 5

A

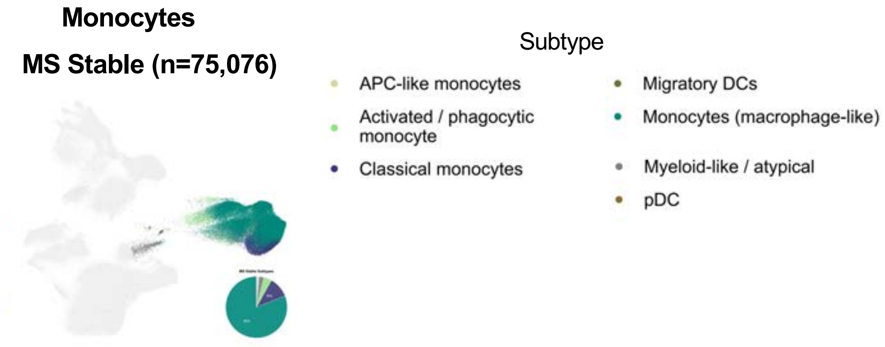


B

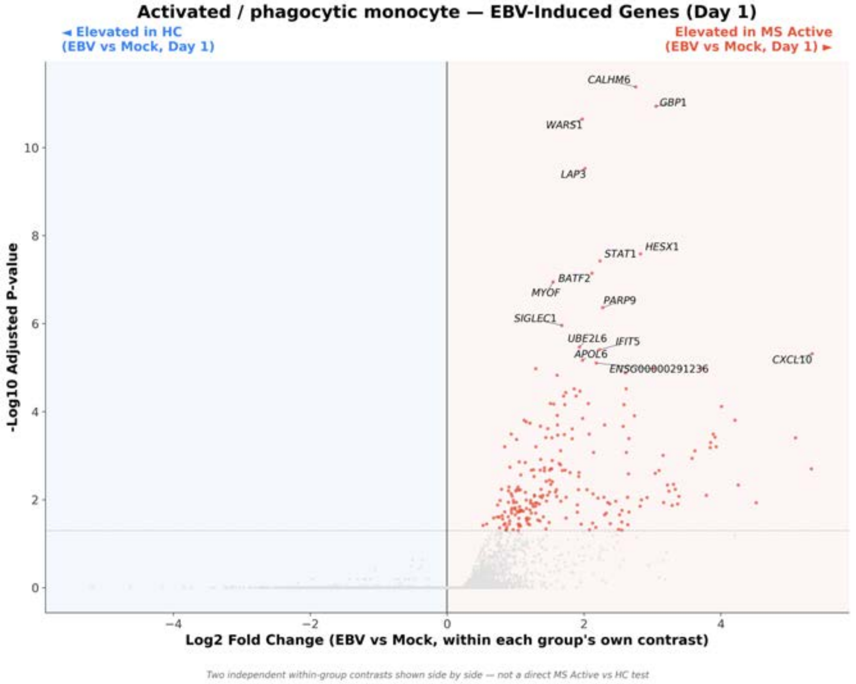


C

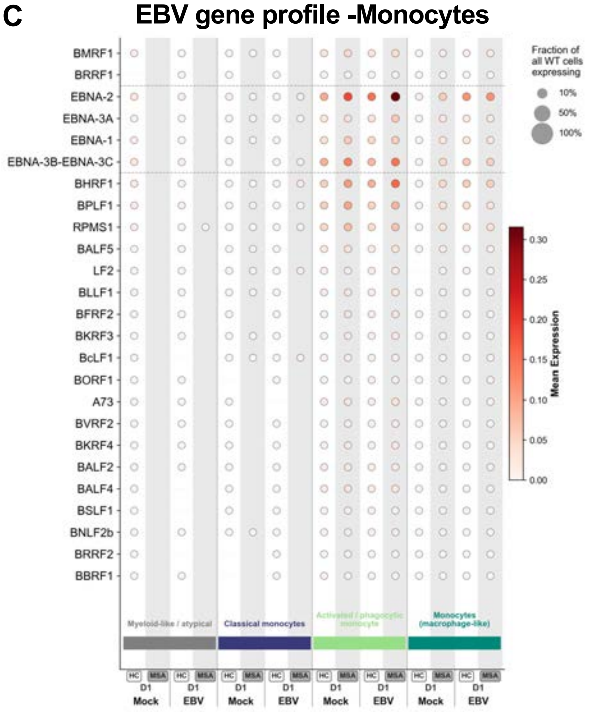

#### Supplementary Figure 6

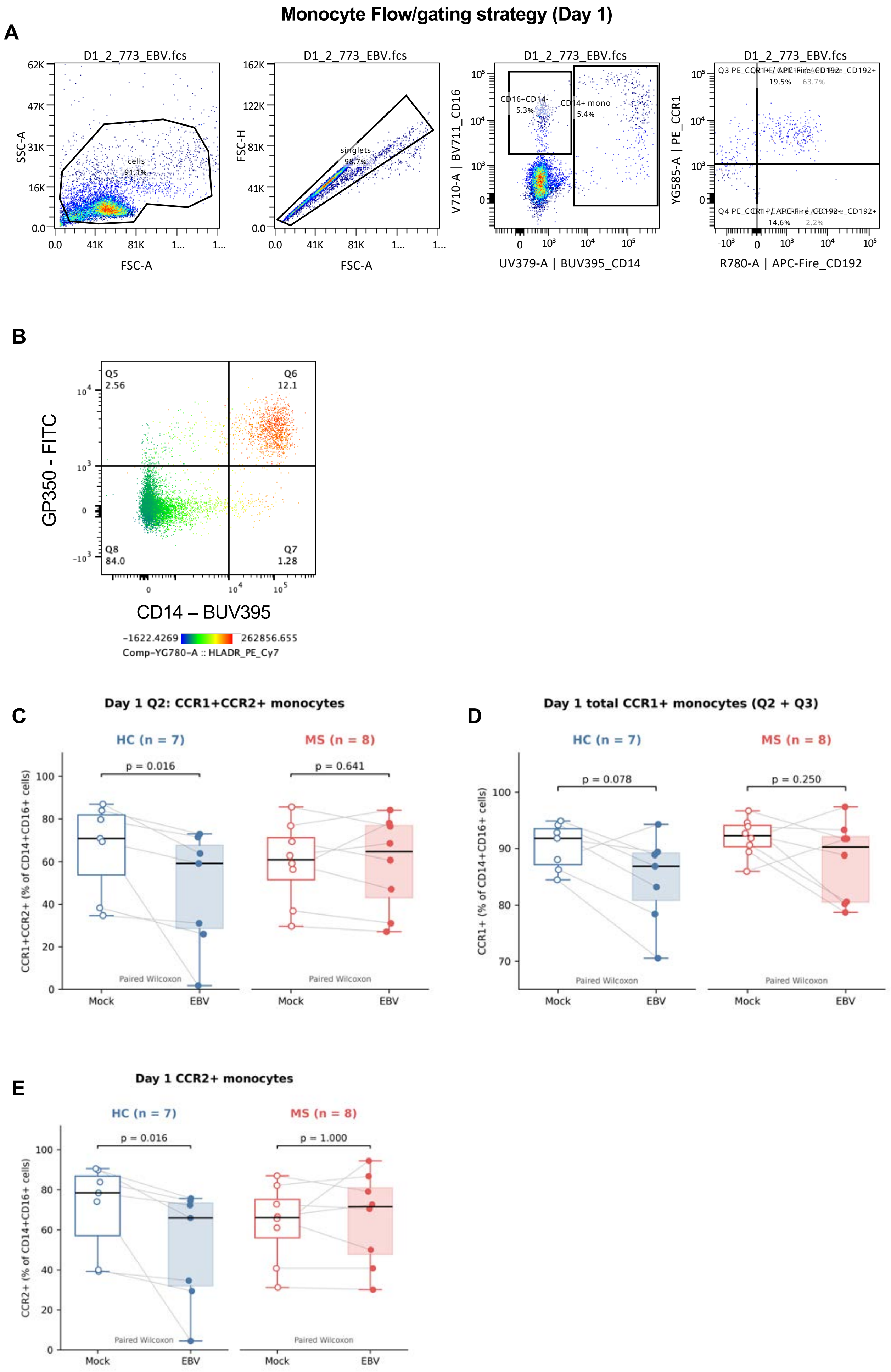

### Supplementary Fig 7

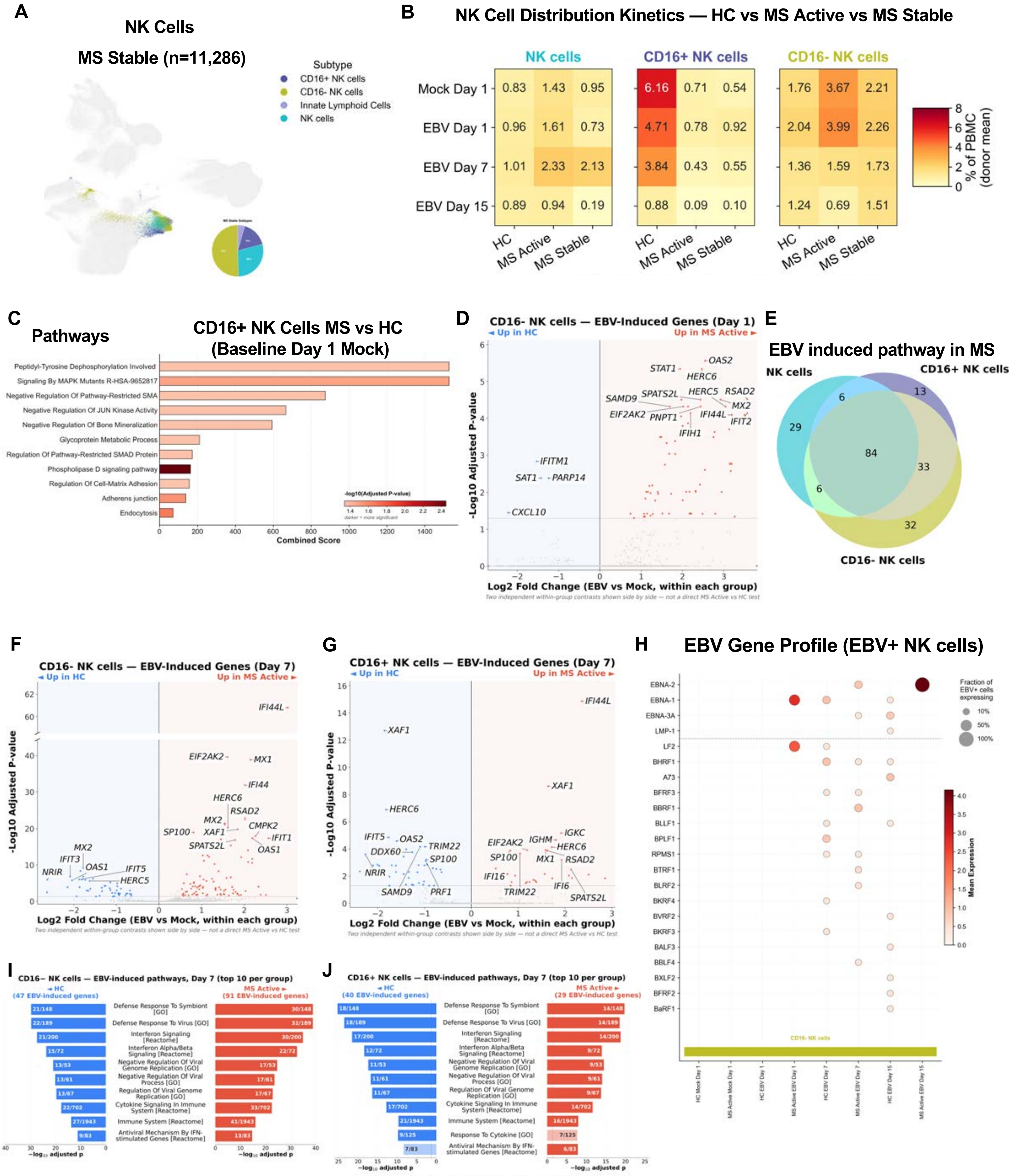

Supplementary Fig 8

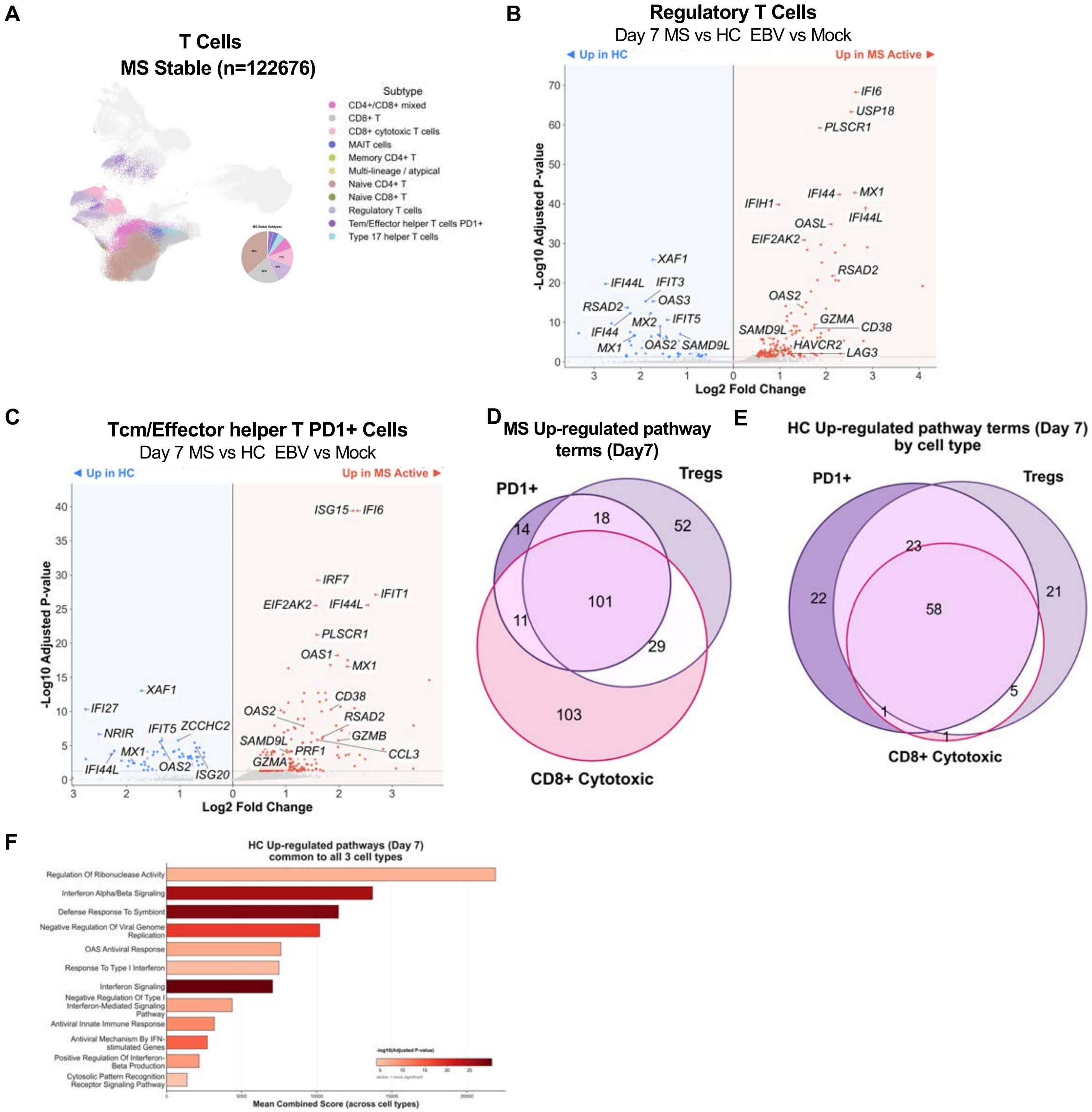

### Supplementary Fig 9

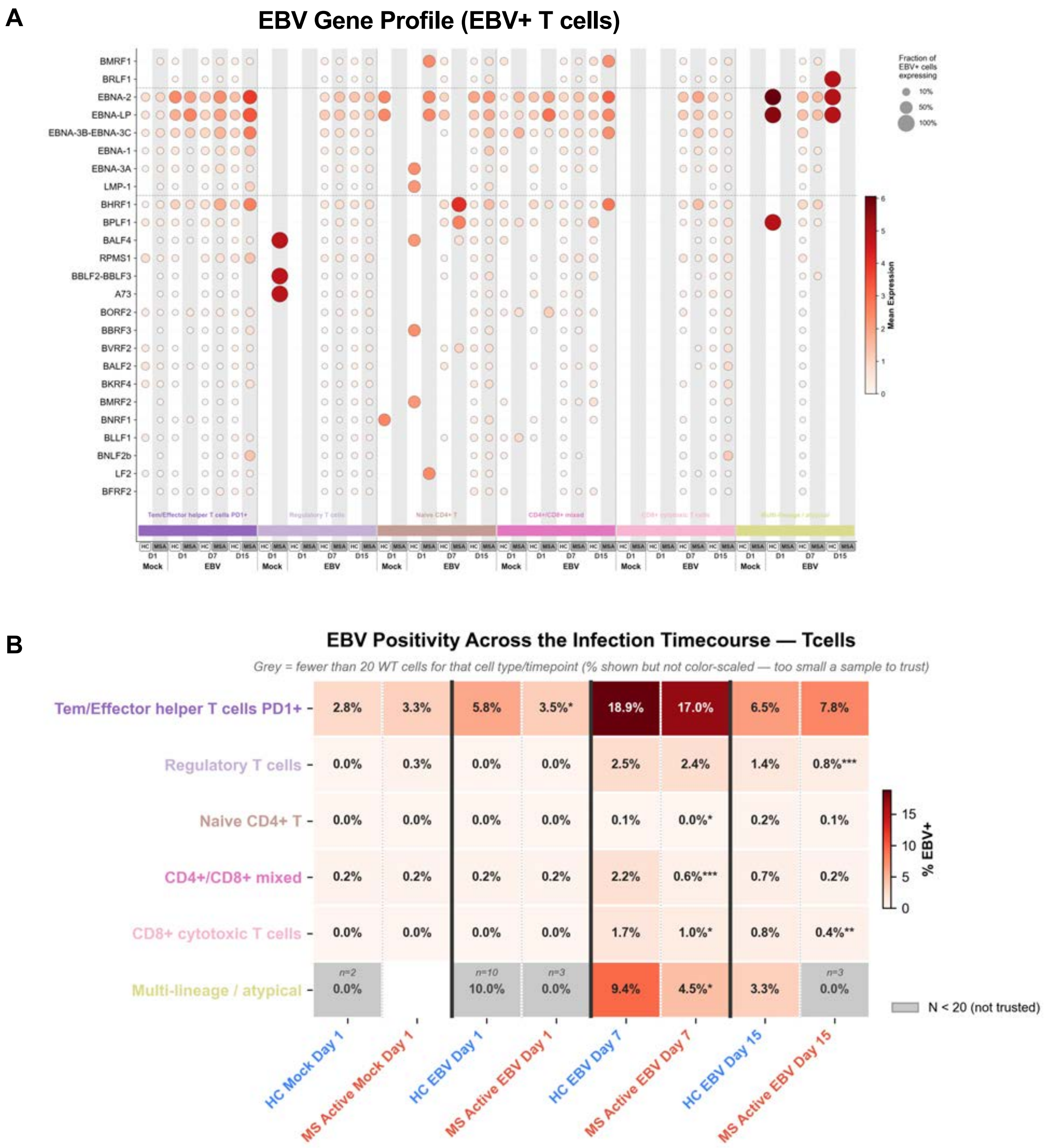

### Supplementary Fig 10

A

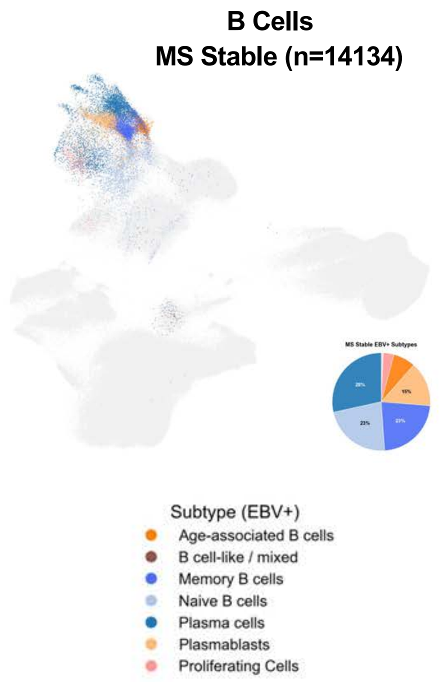


B

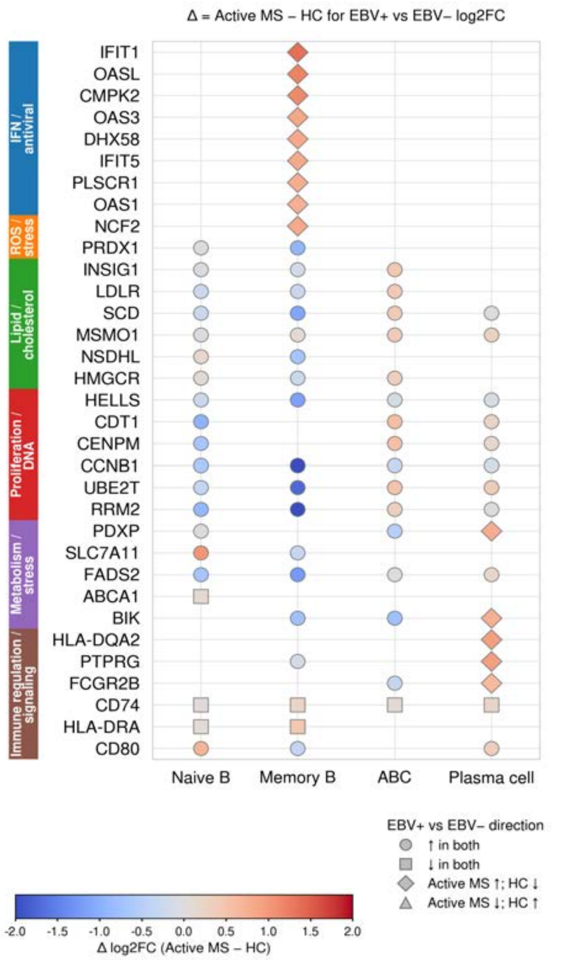


C

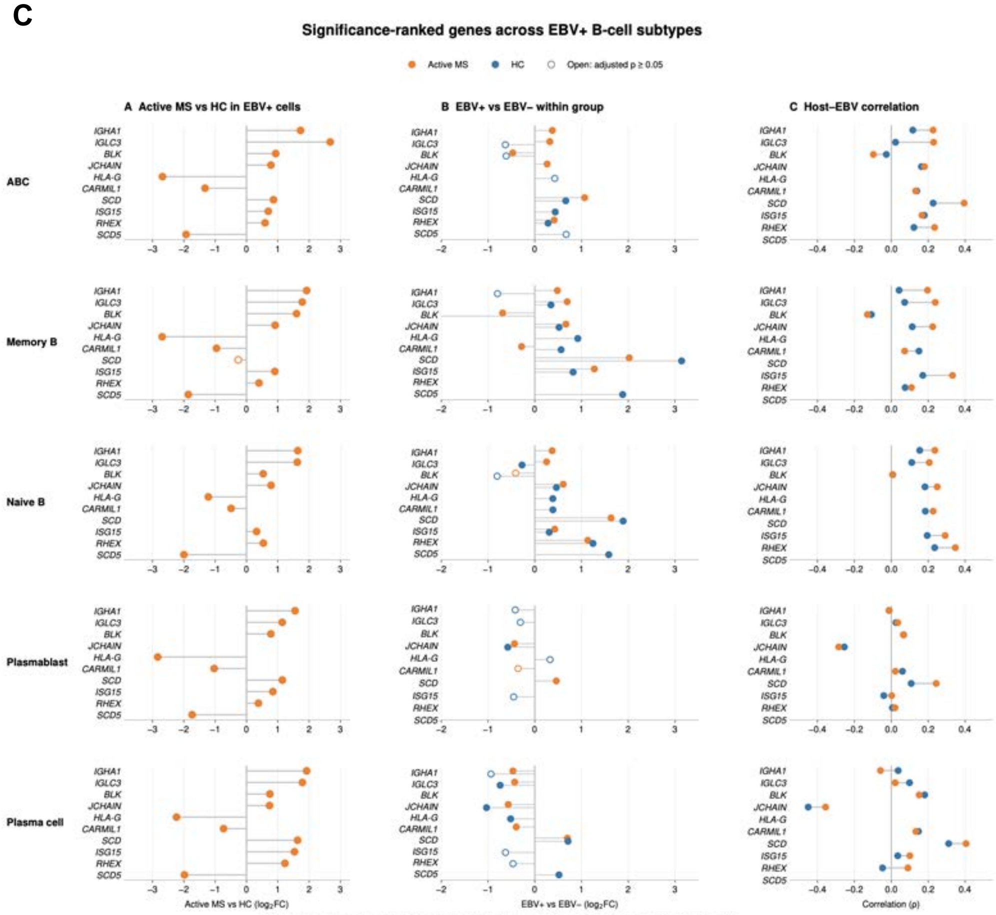


Four requested genes + six significant in all five subtypes (selected by weakest adjusted p). Rows ranked by median −log₁₀(adjusted p).
Missing DE results count as adjusted p = 1 in ranking. Blank = unreported; correlation significance was not supplied.

### Supplementary Fig 11

**A**

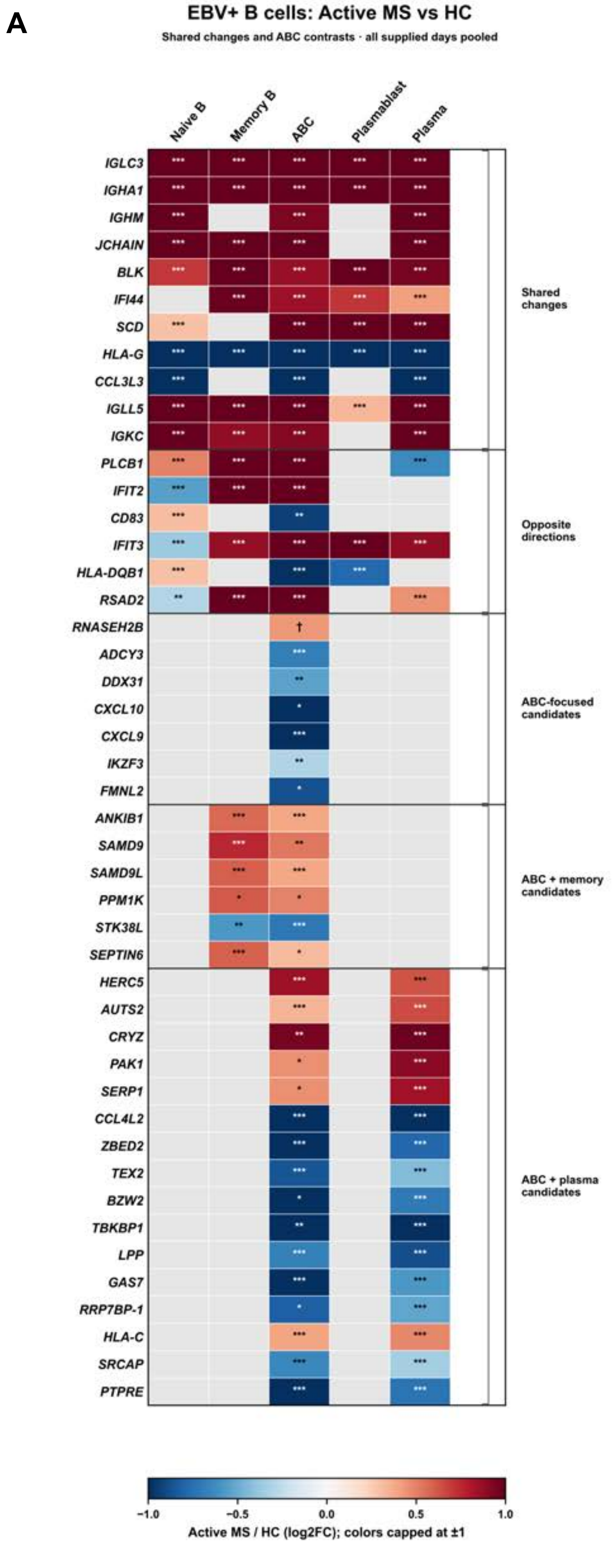


Adjusted P: * <0.05; ** <0.01; *** <0.001 (Bonferroni).
† RNASEH2B: raw P <0.05; adjusted P ≥0.05.
Gray: adjusted P ≥0.05 or absent from the filtered DE table.

**B**

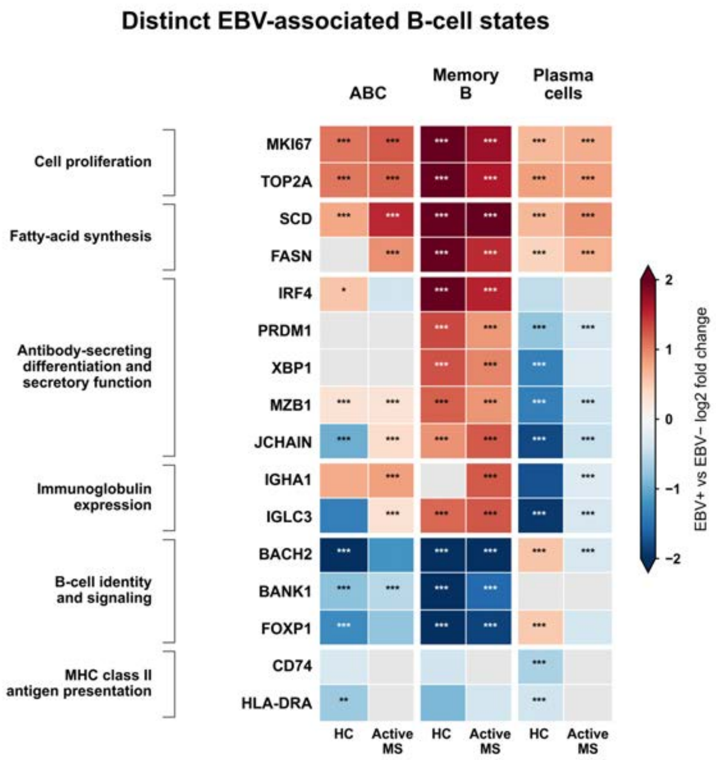


Adjusted P (Bonferroni): * <0.05; ** <0.01; *** <0.001
Gray: absent from the filtered DE table. Colors capped at ±2.
All supplied days pooled; separate EBV+ vs EBV− contrasts within each disease group.

Supplementary Fig 12

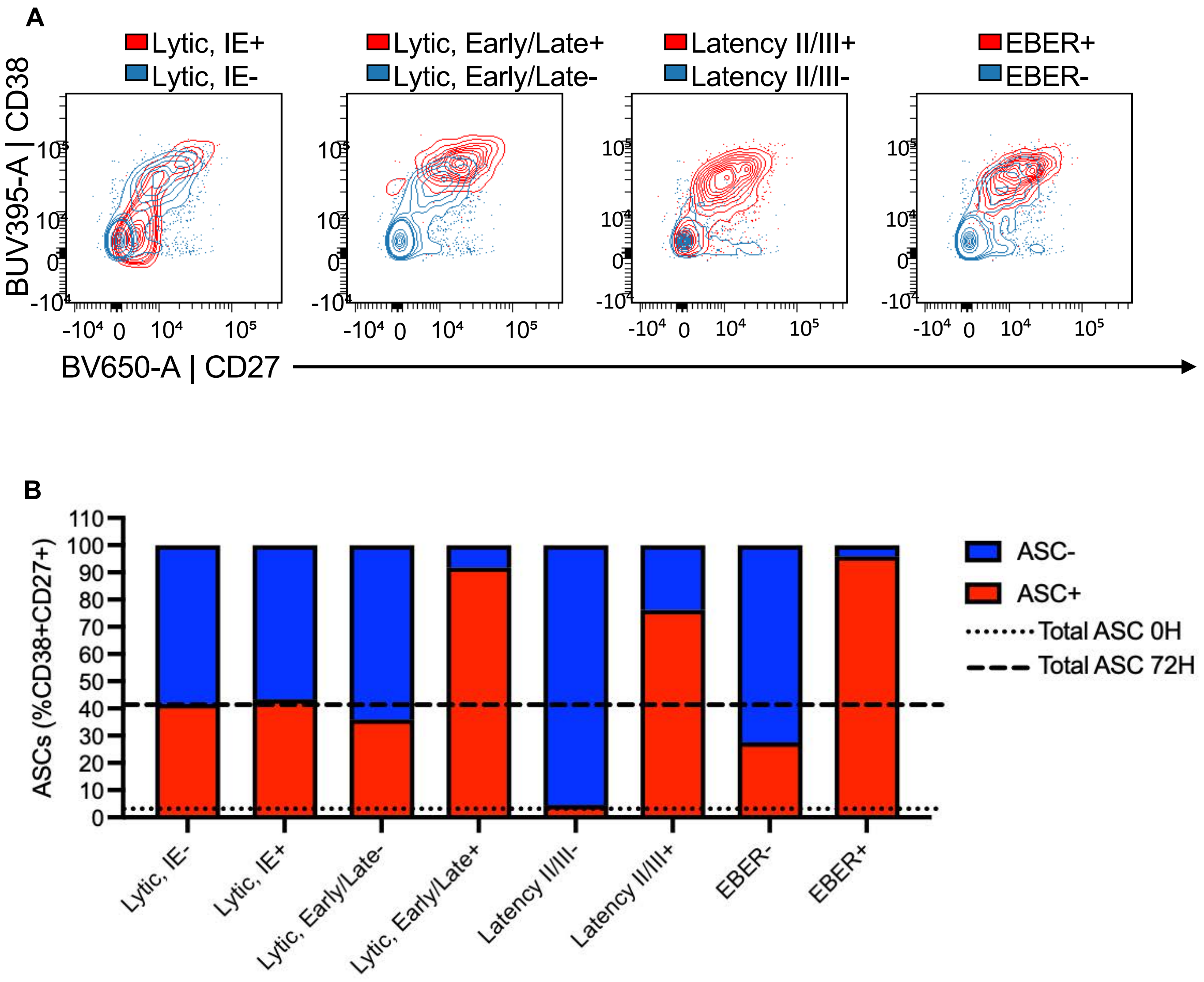

Supplementary Fig 13

A

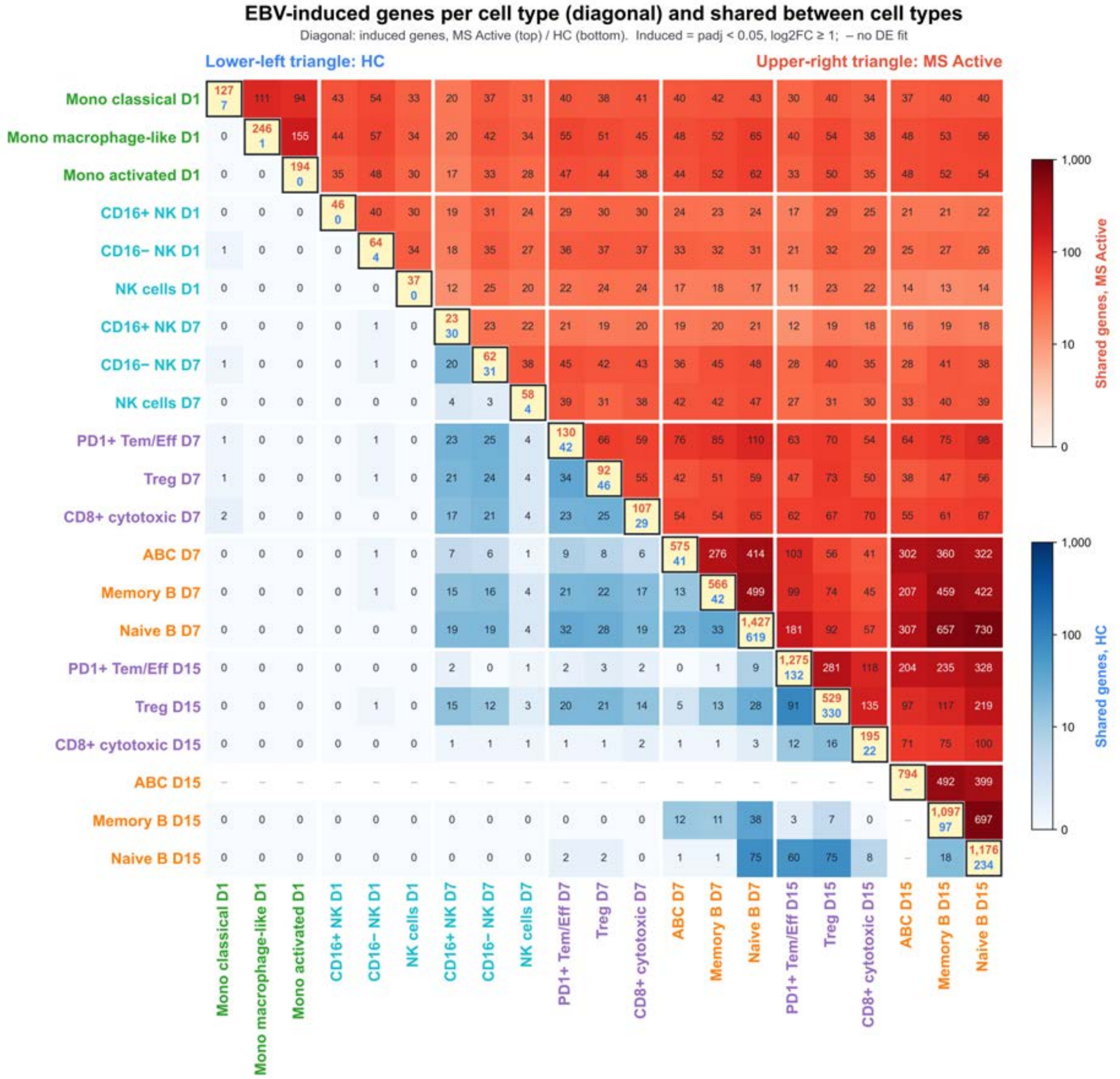


B

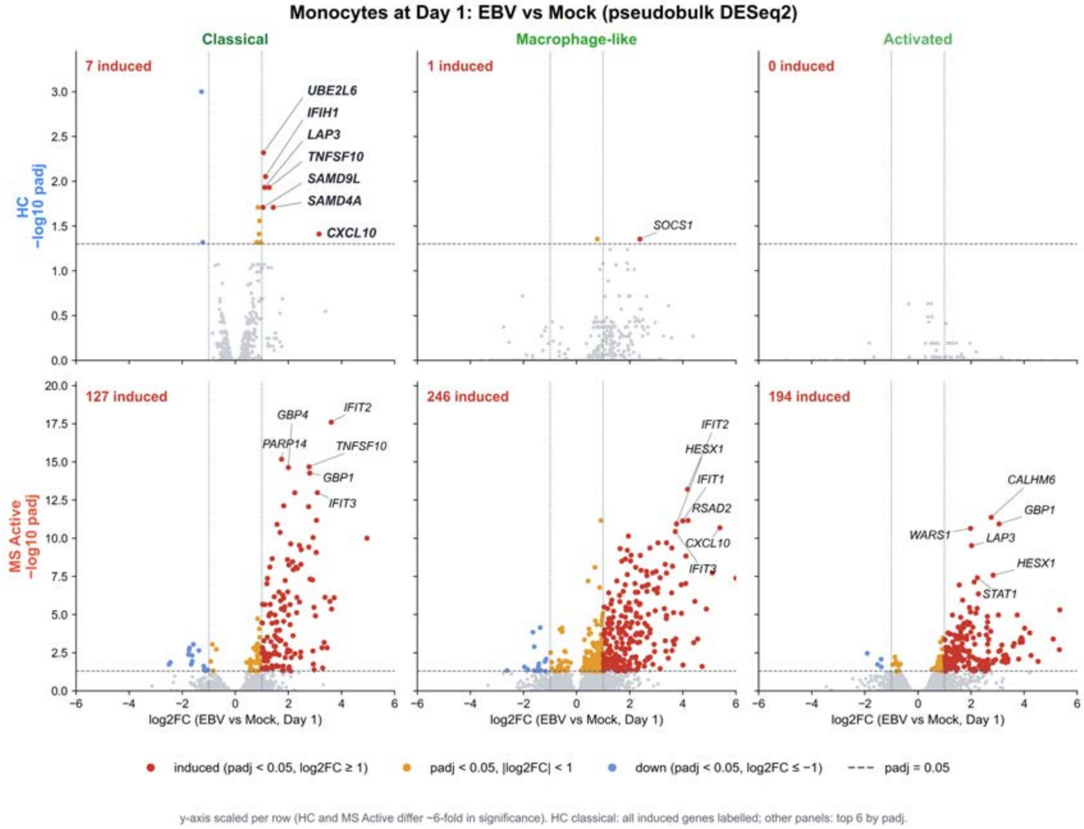

### Supplementary Fig S14

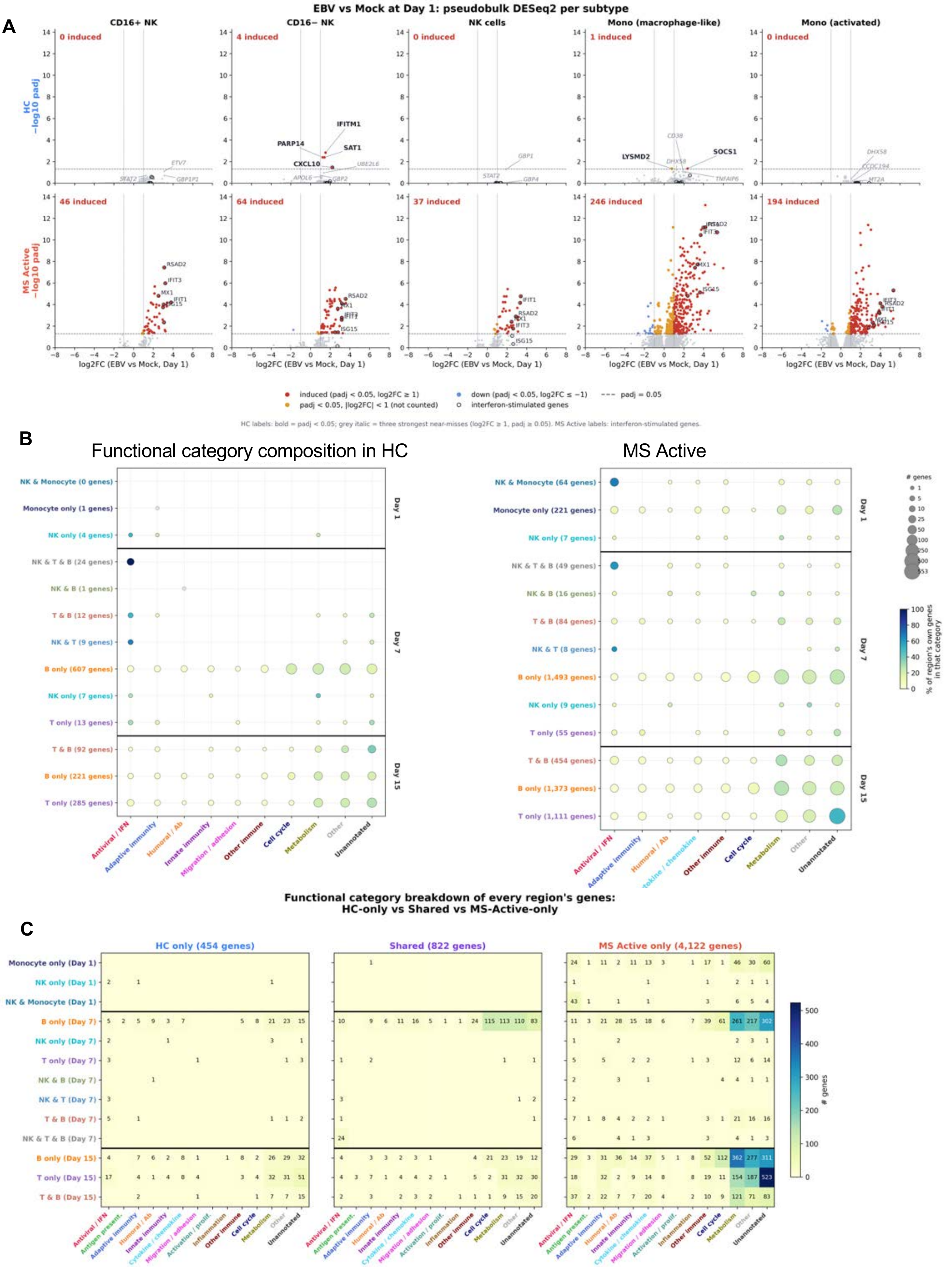

### Supplementary Fig S15

**A**

**EBV Positive Cell Types (MS Active vs HC)**

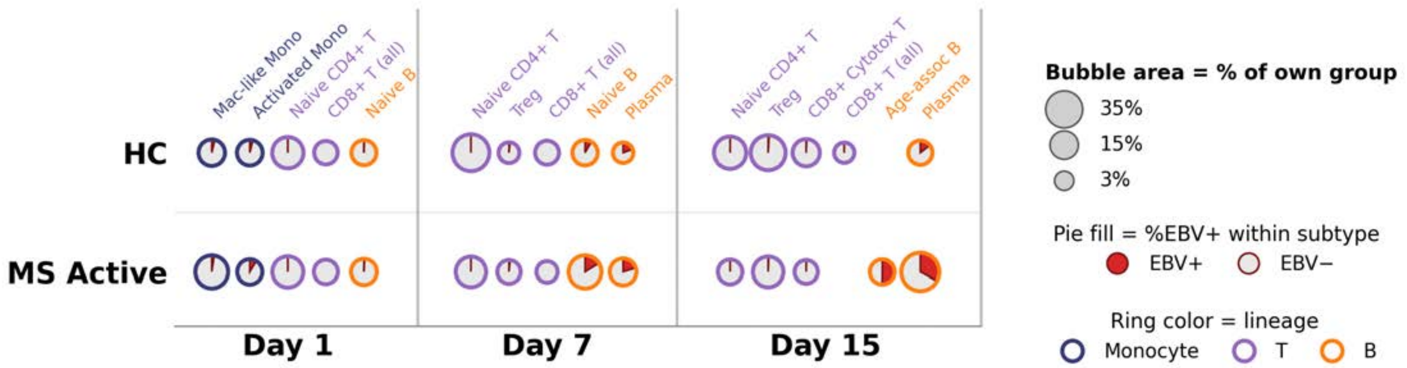


**MS-EBV Gene Signature**

**B**

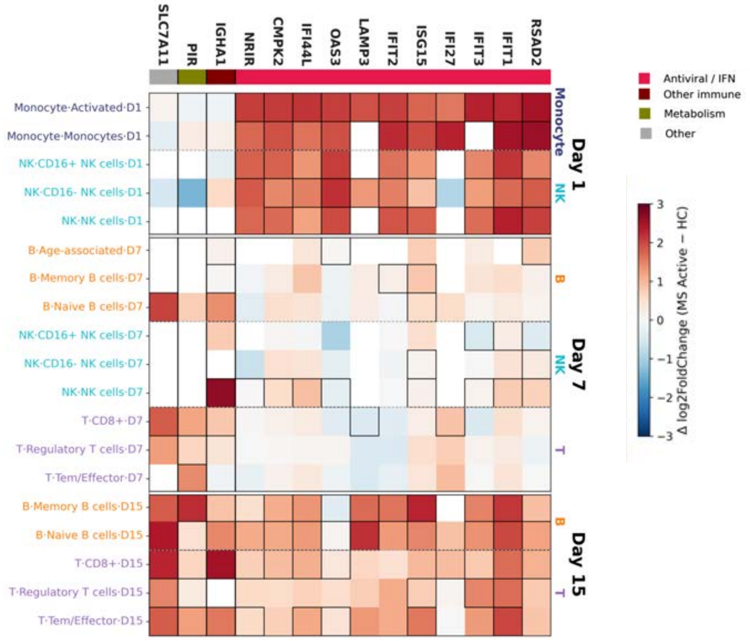
