## Supplementary table1 for "A Dysregulated Peripheral Immune Landscape Favors the Expansion of Epstein-Barr Virus-positive Age-Associated B Cells in Multiple Sclerosis"

### Supplementary Table. Donor cohort and assay participation

| Characteristic | Healthy controls | MS Active | MS Stable | Non-MS neurological controls |
|---|---|---|---|---|
| Donors, n | 15 | 12 | 7 | 5 |
| Age, median (years) | 38 | 34 | 38 | 34 |
| Age, range (years) | 20–50 | 24–51 | 27–59 | 32–72 |
| Female, n | 11 | 7 | 5 | 2 |
| Male, n | 4 | 5 | 2 | 3 |
| Bulk RNA-seq, n | 5 | 7 | 5 | 5 |
| Single-cell RNA-seq, n | 4 | 5 | 3 | 0 |
| Day 1 $CD14^{+}gp350^{+}$ flow cytometry, n | 5 | 3 | 1 | 0 |
| Evaluable for LCL formation, n | 15 | 11 | 6 | 5 |
| LCL formed, n | 7 | 10 | 4 | 2 |
| LCL did not form, n | 8 | 1 | 2 | 3 |
| LCL formation not evaluable, n | 0 | 1 | 1 | 0 |
| Anti-EBNA seropositive at thaw, n/N | 13/13 | 5/5 | 5/5 | 5/5 |

MS Active denotes MS with MRI activity. One donor in the MS Stable group has primary progressive multiple sclerosis (PPMS).

Assay counts are based on recorded participation. A donor may contribute to more than one assay.

LCL, lymphoblastoid cell line. Two donors with low starting cell counts were excluded from the LCL denominator.

Anti-EBNA serology at thaw is reported as positive donors / donors with a recorded result. EBNA, Epstein–Barr nuclear antigen.

Source donor IDs link the two donor-level sheets. Demographics are reported in aggregate.

#### Clinical characteristics by cohort

| Characteristic | Healthy controls | MS Active | MS Stable | Non-MS neurological controls |
|---|---|---|---|---|
| EDSS score, median (n) | 0.0 (n=1) | 2.0 (n=7) | 2.0 (n=7) | 3.5 (n=2) |
| Recent clinical activity, n/N | 3/3 | 12/12 | 1/7 | 2/4 |
| CSF IgG index, median (n) | 0.56 (n=4) | 0.95 (n=12) | 0.77 (n=7) | 0.61 (n=3) |
| CSF oligoclonal bands, median (n) | 0.0 (n=4) | 12.0 (n=12) | 10.0 (n=7) | 0.0 (n=3) |
| CSF white blood cells, median (n) | 1.5 (n=4) | 3.0 (n=7) | 4.0 (n=7) | 3.0 (n=4) |
| CSF protein, median (n) | 30.0 (n=4) | 35.0 (n=7) | 45.0 (n=7) | 41.0 (n=5) |
| Serum NfL z score, median (n) | 0.50 (n=4) | 1.60 (n=5) | 0.81 (n=7) | -0.95 (n=5) |
| Follow-up EDSS score, median (n) | — | 1.2 (n=6) | 2.5 (n=7) | 3.2 (n=2) |

n is the number with a recorded value. n/N is the number with a positive result over the number with data. — indicates no data.

Recent clinical activity covers the prior 6 months. It is recorded separately from MRI-based MS Active status.

CSF white blood cells are in cells/µL and CSF protein is in mg/dL. NfL z scores are age-adjusted.

CSF, cerebrospinal fluid. EDSS, Expanded Disability Status Scale. NfL, neurofilament light chain.

#### Donor assay participation

| Donor ID | Clinical group | Bulk RNA-seq | Single-cell RNA-seq | Day 1 $CD14^{+}gp350^{+}$ flow cytometry | LCL outcome |
|---|---|---|---|---|---|
| 111 | MS Active | Yes | No | No | Not evaluable |
| 120 | Healthy controls | Yes | No | No | Did not form |
| 136 | MS Stable | Yes | No | No | Not evaluable |
| 159 | MS Stable | Yes | No | No | Did not form |
| 160 | Non-MS neurological controls | Yes | No | No | Did not form |
| 187 | MS Stable | Yes | Yes | No | Did not form |
| 96 | MS Active | Yes | No | No | Formed |
| 110 | Non-MS neurological controls | Yes | No | No | Did not form |
| 134 | Non-MS neurological controls | Yes | No | No | Did not form |
| 90 | MS Active | Yes | No | No | Formed |
| 93 | MS Active | Yes | No | No | Formed |
| 107 | MS Active | Yes | No | No | Formed |
| 108 | MS Active | Yes | No | No | Formed |
| 113 | MS Active | Yes | No | No | Formed |
| 192 | MS Stable | Yes | No | No | Formed |
| 132 | Healthy controls | Yes | No | No | Formed |
| 178 | Non-MS neurological controls | Yes | No | No | Formed |
| 179 | Healthy controls | Yes | No | No | Formed |
| 180 | MS Stable | Yes | No | No | Formed |
| 198 | Non-MS neurological controls | Yes | No | No | Formed |
| 246 | MS Active | No | Yes | No | Did not form |
| 248 | MS Active | No | Yes | No | Formed |
| 261 | MS Active | No | Yes | Yes | Formed |
| 263 | MS Active | No | Yes | Yes | Formed |
| 272 | MS Active | No | Yes | Yes | Formed |
| 209 | MS Stable | No | Yes | Yes | Formed |
| 210 | MS Stable | No | Yes | No | Formed |
| 711 | Healthy controls | Yes | No | No | Formed |
| 737 | Healthy controls | Yes | No | No | Formed |
| 728 | Healthy controls | No | Yes | Yes | Formed |
| 686 | Healthy controls | No | Yes | Yes | Did not form |
| 235 | Healthy controls | No | Yes | Yes | Did not form |
| 773 | Healthy controls | No | Yes | Yes | Did not form |
| 783 | Healthy controls | No | No | No | Did not form |
| 514 | Healthy controls | No | No | No | Did not form |
| 763 | Healthy controls | No | No | No | Formed |
| 732 | Healthy controls | No | No | No | Formed |
| 222 | Healthy controls | No | No | Yes | Did not form |
| 730 | Healthy controls | No | No | No | Did not form |

### Clinical characteristics - part 1 of 2

| Donor ID | Clinical group | MS course | EDSS score | Clinical activity within 6 months | Months since most recent symptoms at sampling | CSF IgG index | CSF oligoclonal band count | CSF white blood cells (cells/μL) |
|---|---|---|---|---|---|---|---|---|
| 111 | MS Active | RRMS | 2.0 | Yes | 1.9 | 1.108 | 12 | 4 |
| 120 | Healthy controls | Not applicable | 0.0 | Yes | 8.8 | 0.541 | 0 | 1 |
| 136 | MS Stable | PPMS | 6.0 | No | 28.7 | 0.931 | 7 | 2 |
| 159 | MS Stable | RRMS | 3.0 | No | 31.8 | 0.760 | 12 | 81 |
| 160 | Non-MS neurological controls | Not applicable | 3.0 | Yes | 2.4 | 0.694 | 7 | 4 |
| 187 | MS Stable | RRMS | 0.0 | No | 66.8 | 0.661 | 9 | 0 |
| 96 | MS Active | RRMS | 2.0 | Yes | 2.2 | 0.707 | 10 | 51 |
| 110 | Non-MS neurological controls | Not applicable | | Yes | 7.7 | 0.609 | 0 | 2 |
| 134 | Non-MS neurological controls | Not applicable | | No | | | | 10 |
| 90 | MS Active | RRMS | 2.0 | Yes | 2.8 | 1.105 | 13 | 7 |
| 93 | MS Active | RRMS | 2.0 | Yes | 2.8 | 0.702 | 12 | 0 |
| 107 | MS Active | RRMS | 1.0 | Yes | 3.6 | 0.672 | 5 | 0 |
| 108 | MS Active | RRMS | 2.0 | Yes | 2.6 | 1.037 | 12 | 0 |
| 113 | MS Active | RRMS | 1.0 | Yes | 3.3 | 1.406 | 10 | 3 |
| 192 | MS Stable | RRMS | 0.0 | No | 21.6 | 0.661 | 10 | 27 |
| 132 | Healthy controls | Not applicable | | Yes | 1.4 | 0.577 | 0 | 0 |
| 178 | Non-MS neurological controls | Not applicable | | No | 81.6 | | | 2 |
| 179 | Healthy controls | Not applicable | | Yes | 3.4 | 0.481 | 0 | 9 |
| 180 | MS Stable | RRMS | 1.0 | No | 130.4 | 0.768 | 10 | 2 |
| 198 | Non-MS neurological controls | Not applicable | 4.0 | | 3.6 | 0.447 | 0 | |
| 246 | MS Active | RRMS | | Yes | | 0.560 | 2 | |
| 248 | MS Active | RRMS | | Yes | | 1.570 | 14 | |
| 261 | MS Active | RRMS | | Yes | | 0.870 | 14 | |
| 263 | MS Active | RRMS | | Yes | | 0.730 | 3 | |
| 272 | MS Active | RRMS | | Yes | | 1.340 | 14 | |
| 209 | MS Stable | RRMS | 2.5 | Yes | 1.0 | 0.960 | 10 | 4 |
| 210 | MS Stable | RRMS | 2.0 | No | 3.0 | 1.412 | 18 | 9 |
| 711 | Healthy controls | Not applicable | | | | | | |
| 737 | Healthy controls | Not applicable | | | | | | |
| 728 | Healthy controls | Not applicable | | | | | | |
| 686 | Healthy controls | Not applicable | | | | | | |
| 235 | Healthy controls | Not applicable | | | | | | |
| 773 | Healthy controls | Not applicable | | | | | | |
| 783 | Healthy controls | Not applicable | | | | | | |
| 514 | Healthy controls | Not applicable | | | | | | |
| 763 | Healthy controls | Not applicable | | | | | | |
| 732 | Healthy controls | Not applicable | | | | | | |
| 222 | Healthy controls | Not applicable | | | | 0.616 | 0 | 2 |
| 730 | Healthy controls | Not applicable | | | | | | |

RRMS, relapsing-remitting multiple sclerosis. PPMS, primary progressive multiple sclerosis.

DMT, disease-modifying therapy.

### Clinical characteristics - part 2 of 2

| Donor ID | Clinical group | CSF protein (mg/dL) | Age-adjusted serum NfL z score | Follow-up (years) | MRI activity during follow-up | Follow-up EDSS score | Steroids within 4 weeks | DMT history |
|---|---|---|---|---|---|---|---|---|
| 111 | MS Active | 16 | | 2.21 | No | 1.5 | | |
| 120 | Healthy controls | 22 | 0.36 | | | | | |
| 136 | MS Stable | 55 | 1.85 | 1.93 | No | 6.5 | | |
| 159 | MS Stable | 53 | -0.33 | 1.93 | No | 3.0 | | |
| 160 | Non-MS neurological controls | 26 | 2.01 | 1.36 | No | 2.5 | | |
| 187 | MS Stable | 50 | 0.81 | 1.77 | Yes | 2.0 | No | 0 |
| 96 | MS Active | 35 | 1.08 | 2.42 | No | 1.0 | | |
| 110 | Non-MS neurological controls | 63 | -0.95 | | | | | |
| 134 | Non-MS neurological controls | 41 | 0.33 | | | | | |
| 90 | MS Active | 29 | 2.51 | 2.41 | No | 2.0 | | |
| 93 | MS Active | 61 | | 2.41 | No | 1.5 | | |
| 107 | MS Active | 47 | 2.65 | 1.55 | No | 0.0 | | |
| 108 | MS Active | 24 | 1.23 | 0.90 | No | | | |
| 113 | MS Active | 38 | 1.60 | 1.85 | No | 1.0 | | |
| 192 | MS Stable | 45 | 0.23 | 1.37 | Yes | 0.0 | | |
| 132 | Healthy controls | 45 | 3.19 | | | | | |
| 178 | Non-MS neurological controls | 27 | -2.99 | | | | | |
| 179 | Healthy controls | 38 | 0.64 | | | | | |
| 180 | MS Stable | 33 | -1.23 | 1.84 | No | 1.0 | | |
| 198 | Non-MS neurological controls | 59 | -1.60 | 1.86 | Yes | 4.0 | | |
| 246 | MS Active | | | | | | No | 0 |
| 248 | MS Active | | | | | | No | 0 |
| 261 | MS Active | | | | | | No | 0 |
| 263 | MS Active | | | | | | No | 0 |
| 272 | MS Active | | | | | | No | 0 |
| 209 | MS Stable | 28 | 1.98 | 1.68 | Yes | 3.0 | No | 0 |
| 210 | MS Stable | 31 | 3.29 | 1.07 | Yes | 2.5 | No | 0 |
| 711 | Healthy controls | | | | | | | |
| 737 | Healthy controls | | | | | | | |
| 728 | Healthy controls | | | | | | | |
| 686 | Healthy controls | | | | | | | |
| 235 | Healthy controls | | | | | | | |
| 773 | Healthy controls | | | | | | | |
| 783 | Healthy controls | | | | | | | |
| 514 | Healthy controls | | | | | | | |
| 763 | Healthy controls | | | | | | | |
| 732 | Healthy controls | | | | | | | |
| 222 | Healthy controls | 20 | -1.60 | | | | | |
| 730 | Healthy controls | | | | | | | |

RRMS, relapsing-remitting multiple sclerosis. PPMS, primary progressive multiple sclerosis.

DMT, disease-modifying therapy.
